# How learned expectations shape brain-wide responses

**DOI:** 10.64898/2025.12.15.694430

**Authors:** Ari Liu, Michael Schartner, International Brain Laboratory, Ila Fiete

**Affiliations:** The International Brain Laboratory; Massachusetts Institute of Technology, Cambridge, MA, USA; Champalimaud Foundation, Lisbon, Portugal

## Abstract

Expectations stemming from prior knowledge have long been known to influence stimulus processing and decision-making. Empowered by brain-wide recordings during a two-choice sensory decision-making task from the International Brain Laboratory^1^, we sought to identify the form of expectation-based modulations and how they influence ongoing computations. To discern where, when, and how prior knowledge about the stimulus – known to be stored in the brain over the inter-trial period^2^ – influences brain activity over the arc from stimulus processing to choice selection and action generation, we disentangled the encoding of the highly correlated stimulus, choice, and prior variables as sensory processors, choice/integration processors, and movement generators. We examined which brain regions were modulated by prior expectations and at which point in the task, and characterized whether those modulations take the form of biasing initial activations or response gains. We found that the activities of choice/integrators and movement generators were influenced by prior expectation, while purely stimulus processors were not significantly modulated by prior expectation. In addition, we found that expectation-based modulation takes the form of both initial activity bias and gain modulation in integration/choice computations and in movement generators. We further find that the brain-wide recordings reveal an emergent simplicity: A few-parameter mechanistic neural circuit model captures the population dynamics of stimulus response, integration/choice processing, and action initiation, as well as their modulations by the prior. After fitting to physiology, the model’s behavior predicts the psychometric and chronometric behavior of mice with no additional parameters. It reveals that the larger bias and gain modulations of movement initiators relative to integration/choice in the data and model is inherited by amplifying integrator/choice inputs rather than being an additional expectation-based effect. Collectively, our results characterize in fine spatiotemporal detail and brain-wide breadth the mechanisms and loci where prior expectations influence cognition, while the novel synthesis between large-scale electrophysiology and mechanistic modeling reveals an organizing simplicity behind perceptual decision making across the brain.

## Introduction

Using past experience to modify future behavior is a hallmark of cognitive decision making. In stochastic settings, prior knowledge usually takes the form of learned estimates of input-response-outcome probabilities, or *priors*, and using this knowledge to shape responses. Behaviorally, various animals from flies ^3^ to humans ^2,4,5^ exhibit such learning and use it to modulate their decisions. Prior knowledge thus influences responses to inputs, but its locus and mechanisms of action – when, where, and how this modulation enters along the processing pathway from input to response – requires elucidation.

The question of loci – whether early stimulus processing or only later stages of choice formation and action initiation are modulated by prior expectations – remains open ^6–14^. On how and when in the decision-making process priors shape ongoing processing – whether by adding activity biases at the start of processing, or by altering the gain of inputs and responses during processing, the evidence is mixed ^7,10,11,13–19^. This inconclusive state of understanding can partly be traced to the limited scope of recordings (small numbers of animals, neurons, and brain regions) and the correlated nature of the relevant variables (the stimulus and choice are typically matched in well-trained perceptual decision makers, and by definition the stimulus and prior expectation about the stimulus are correlated in animals correctly estimating the prior). Variations in the tasks used in different studies mean the results cannot be easily combined to draw stronger conclusions.

Here, we consider the question using physiological recordings from the International Brain Laboratory (IBL) ^1^, comprising ∼ 6 × 10^5^ neurons recorded across the brain (208 Beryl regions included in our analysis) in 139 expert-level mice performing a perceptual decision-making task that relies on the incorporation of prior knowledge about the side that the stimulus will likely be presented. Our work builds on comprehensive previous analyses describing brain-wide neural responses during the IBL task ^1^ and identifying where prior expectations are stored between trials ^2^.

In what follows, we seek to determine when, where, and how priors intervene in the arc from sensation to decision to action. We do so by first identifying the decorrelated encoding loci for visual stimulus response, integration/choice, and movement initiation. Then, we examine whether the prior modulates responses at these loci, and if it does, whether it modifies their initial activity biases (biases) or the gains of their inputs. We find that the prior strongly modulates the initial biases of integration/choice and movement initiation responses and the gain of integration/choice responses, but do not find evidence of prior modulation in stimulus processing. We further find that a simple mechanistic circuit model captures the key dynamics of neural activity for sensory decision making and prior modulation, quantifies the strengths of prior modulation, and reveals that movement initiation gain modulation is inherited from integration/choice processing rather than through an additional prior effect.

## Results

In the IBL task, mice determine whether a visual stimulus appeared on the left or right and indicate their decision by turning a wheel, and correct responses yield a reward (**Fig.** 1a). Over a consecutive block of trials, stimuli appear on one side with high probability (80%); the high-probability side then flips without an explicit cue. Mice learn to recognize and exploit this block structure, using it as prior knowledge in generating more accurate responses to faint stimuli, **Fig.** 1b. The high-density and brain-wide recordings of the IBL make it possible to study where, when, and how prior representations in the brain shape decision-making, **Fig.** 1c. We first characterize how and where key task variables are represented in the brain, then examine how these representations are modulated by prior knowledge within each trial.

**Figure 1.**
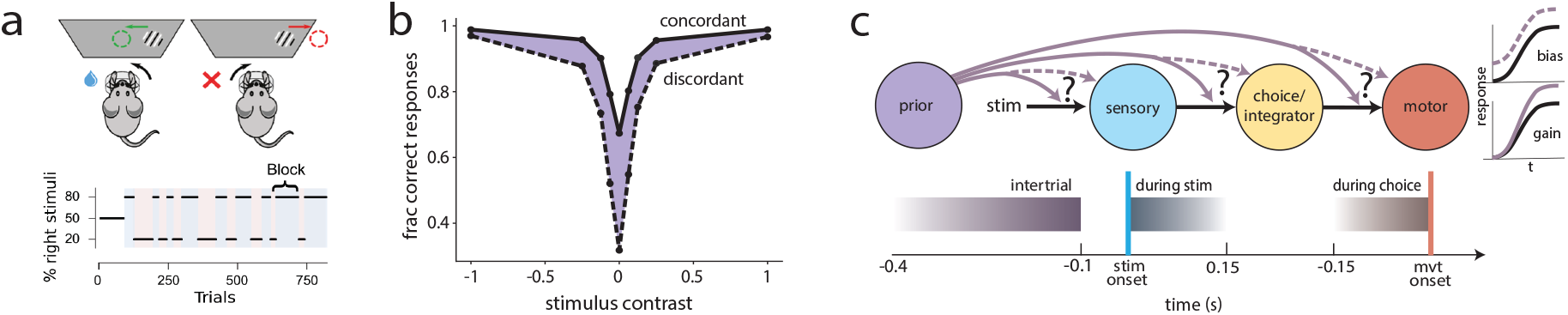
Task, behavior, and questions. **a** Experimental setup ^1^ with example block structure. **b** Trained mice use block information to solve the task, quantified by the gap between the solid (concordant: stimulus on the high-probability block side) and dashed response (discordant or stimulus on opposite of the high-probability block side) curves. The difference is especially visible for zero-contrast trials, for which animals can only use block side information as the stimulus is invisible. **c** Top: The question we would like to address: How does the prior influence ongoing computations during a trial? We consider whether the prior affects response gains or biases (purple solid versus purple dashed), and which stages of the computation it influences. Bottom: Time windows used for analysis.

### Identifying decorrelated stimulus, integration, movement and prior sensitivity

The block (prior) side, the stimulus side, and the choice side comprise the key variables in the IBL task, and their combinations define 8 trial types (e.g., *R* Stimulus, *R* Choice, *L* Block, which we will abbreviate as *S*_*R*_*C*_*R*_*B*_*L*_). Before determining which computations are modulated by the prior, we must grapple with the problem that the variables are highly correlated: on a left block, the stimulus is far more likely to appear on the left side, and in well-performing animals, the choice is also highly likely to be on the left. Similarly, because the stimulus is more likely to appear on the same side as the prior, by definition, the coding of the prior will be highly correlated with coding of the stimulus. To identify specific encoding of stimulus side, choice side, or prior side despite these correlations, we used explicit conditioning on the different binary variables as described below.

For instance, to isolate sensitivity to the stimulus, we compared left versus right stimulus trials separately within each of the four block, choice dichotomies (e.g. comparing left versus right stimulus responses within the right choice, right block condition). Within each dichotomy, we performed population trajectory analyses as described in IBL’s brain-wide map paper ^1^.

Briefly, we combined all neurons across animals in a single “supersession” as follows: for each neuron *α*, we aggregated its responses within each of the eight stimulus, choice, prior dichotomies, aligned the aggregated responses to a trial landmark (e.g. stimulus onset or movement onset), and computed the time-varying average response 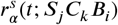 across trials, aligned to stimulus onset (superscript *s*; movement onset alignment would use the superscript *m*) in the block side *B*_*i*_, stimulus side *S*_*j*_, and choice side *C*_*k*_ condition. The stimulus-aligned responses consist of an interval of 150 ms from the stimulus onset, while movement-aligned responses consist of the time interval of 150 ms before movement onset. For stimulus responses, we further define an “early” stimulus response period (superscript *se*), consisting of the interval [0,80] ms from stimulus onset. Informed by ^1^, transient sensory responses are better captured during this shorter interval, whereas the longer (150 ms) stimulus aligned time window is good for identifying stimulus integration/choice signals.

To determine sensitivity to stimulus side, decorrelated from sensitivity to choice and prior side, we computed the Euclidean distance between the left and right stimulus responses within each block and choice condition. By fixing block and choice sides as the stimulus side response difference is computed, the effects of correlated variation are removed. The sum of these differences across all four block, choice conditions for cell *α* yields a total L, R stimulus response difference curve 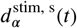. We can compute these differences for cell populations by replacing the neuron response 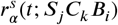 by a neural response vector 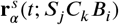, where *α* designates the cell population and **r** designates the vector of responses of all cells in the designated population, with L,R stimulus difference and choice difference curves based on the distance between response trajectories computed in an *N*_*α*_-dimensional space, where *N*_*α*_ designates the number of neurons in group *α*:

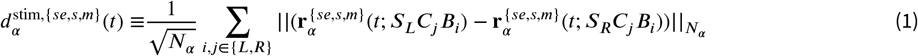

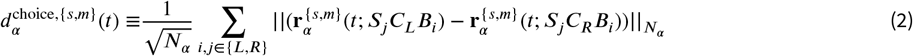

where 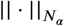 corresponds to the Euclidean distance computed per time-point between the two population trajectories (if *α* designates a cytoarchitecturally defined Beryl region, *N*_*α*_ corresponds to all the neurons in the dataset from that region; if *α* designates a functionally defined set of neurons, then *N*_*α*_ refers to the number of neurons in this set).

We also examine a less strict definition of early stimulus sensitivity by only controlling for prior side (and loosening the control with choice side, assuming choice sensitivity only arises later on in a trial):

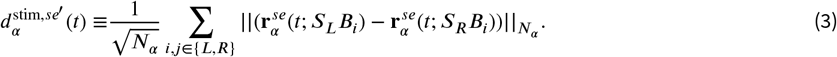

To assess significance in the resulting difference curves, we generate a null distribution of control curves by shuffling labels of the variables of interest. For example, we shuffle left and right labels for the stimulus side within each choice/block dichotomy to generate shuffle controls of 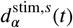, and compare the mean of the difference curve 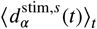 with the distribution of the shuffle controls’ mean values to calculate a p-value, with false discovery rate (FDR) correction at a *p*-value threshold of 0.01 (details in Methods).

This procedure reveals that early stimulus sensitivity is decorrelated with prior in some brain regions (e.g. VISp, VISal) and lack of significant early stimulus selectivity in most regions (e.g. MOs), **Fig.** 2a and **Fig.** S1. When controlling for correlations with both prior and choice, early stimulus sensitivity is only weakly significant (0.01 < *p* < 0.05) in a few regions (e.g. VISal, VISpm), **Fig.** S1, S2 Some regions like VeCB are stimulus sensitive without a pronounced early stimulus response, **Fig.** 2b. Overall, few regions have significant decorrelated stimulus sensitivity, **Fig.** S1. In contrast, we more broadly find decorrelated integration/choice sensitivity in the form of ramping activation in regions like GRN, whether aligned to stimulus or movement onset, **Fig.** 2c. Overall, a large number of regions exhibit high stimulus-aligned or movement-aligned choice sensitivity, **Fig.** S1.

**Figure 2.**
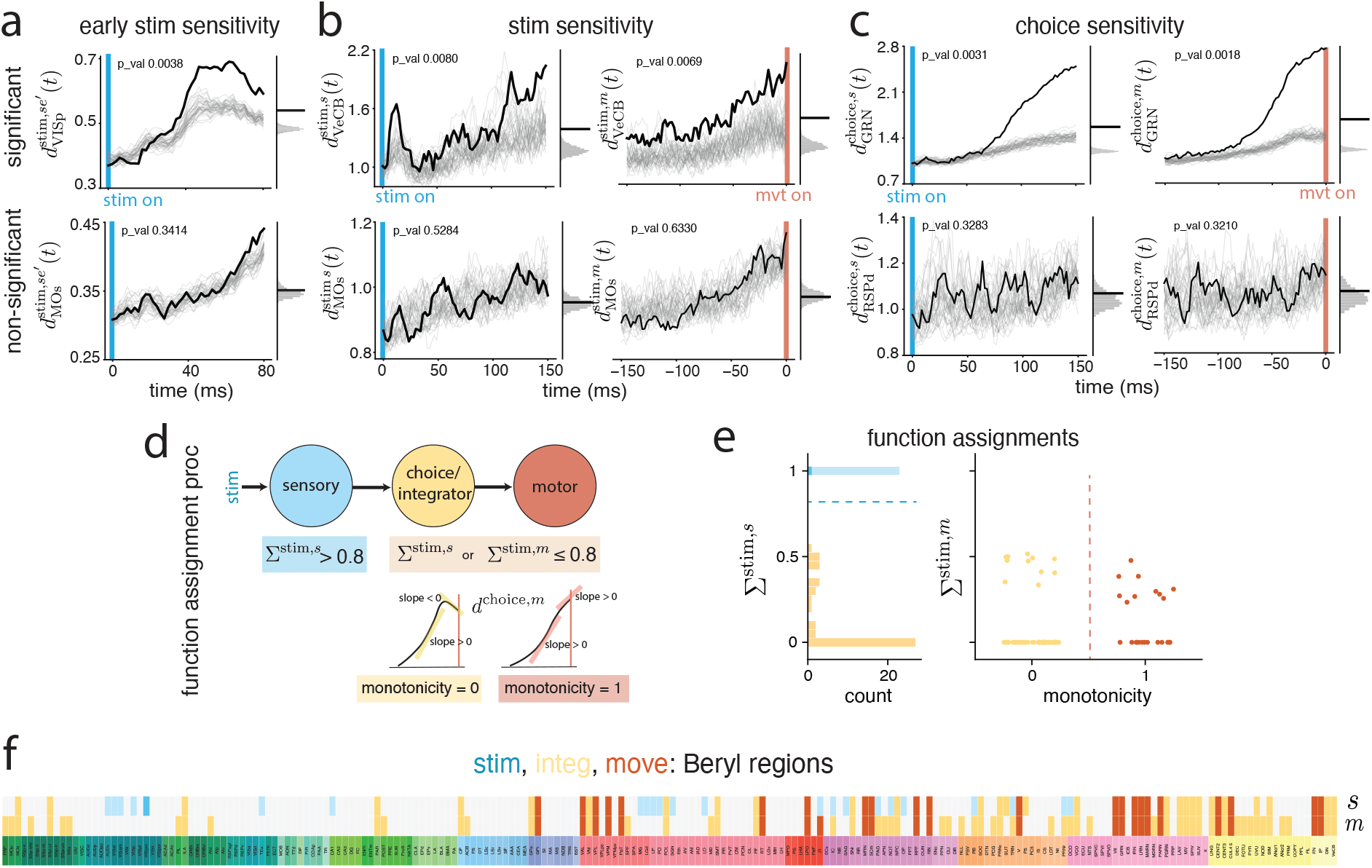
Identification of stimulus side, choice side, and prior side sensitivity and classification of sensation, integrator/choice, and movement initiation areas via control of correlated variables. **a** Early stimulus side sensitivity: euclidean distance of left versus right stimulus response conditioned on block sides, for the first 80 ms after stimulus onset (black curve), with shuffle controls (gray curves: randomize the left and right stimulus labels before subtracting within a condition). Right: distribution of shuffle curve means (gray histogram), to compare with the true data mean (black curve) to generate a *p*-value. Example regions with significant (top, *p* < 0.01) versus non-significant (bottom) stimulus side responses. Example regions for early stimulus side sensitivity conditioned on both block and choice sides are not shown here since no regions are significant with *p* < 0.01, but results with *p* < 0.05 are summarized in **Fig. S2a**. **b** Stimulus side sensitivity conditioned on block and choice sides, 0 − 150 ms after stimulus onset (left) and 0 − 150 ms before movement onset (right). **c** Choice side sensitivity conditioned on block and stimulus sides, aligned to stimulus and movement onset. **d** Procedure for assigning functions to regions: Σ^stim^ differentiates purely stimulus processors from the rest (integration/choice regions and movement initiation areas) and a monotonicity metric differentiates the integrator/choice regions from movement initiation areas (see **Fig. S1** for full table of region-wise function assignments). **e** Functional classification of regions. Left: histogram of Σ^stim,s^, with Σ^stim,s^ > 0.8 defining purely sensation regions. Darker blue: sensory regions defined stringently by controlling for correlations with both choice and prior in early stimulus sensitivity (Σ^stim,s^ > 0.8 using 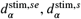), results in only one region; lighter blue: sensory regions defined by controlling only for prior in early stimulus sensitivity (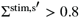 using 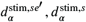); orange: choice/integration or motor regions. Right: Strip plot of Σvs the monotonicity class defines movement initiation areas (red) (Σ≤ 0.8 and monotonicity = 1; dots are Beryl regions, colored by their classification). The rest of the regions with Σ^stim,s^ ≤ 0.8 or Σ^stim,m^ ≤ 0.8 are integrator/choice regions (yellow) when aligned to stimulus or movement onset. **f** Functional classification of all recorded regions into 4 categories (same color scheme as in **e**), one row per window type, stimulus-onset aligned *s* or movement-onset aligned *m*. Gray: no significant stimulus/ early stimulus/ choice sensitivity, thus not categorized to any of the functions.

### Sensory, integration/choice, and movement processing

We can assign a function to each group of neurons *α*, based on this set of decorrelated sensitivity measures for stimulus, choice, and prior. Our goal is to understand how these distinct functions, carried out at different times within a trial, are modulated by the prior. Possible functional assignments are “pure” stimulus processors (with little integration/choice sensitivity; we will call these stimulus processing neurons); integration/choice processors that are not movement initiators (we will call these integrator/choice neurons); and movement initiators.

We reason that stimulus processors should significantly encode the decorrelated stimulus difference, do so early in the stimulus period, and be insensitive to integration/choice relative to integrator/choice processors. To differentiate stimulus processing from integration/choice processing, we combine the sensitivity metrics into a classification axis, Σ^stim,*s*^, which quantifies the relative strength of stimulus sensitivity versus integration/choice sensitivity in stimulus-aligned responses (**Fig**.S1a):

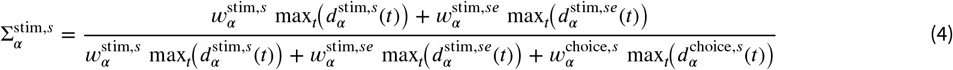

where 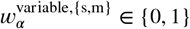 indicates statistical significance for 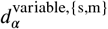 (details in Methods). This metric is only defined for neuron groups with significant stimulus or choice coding (106/208 Beryl regions), and then it takes values Σ^stim,*s*^ ∈ [0, 1]. Σ^stim,*s*^, with high values for neurons that have strong early stimulus side or overall stimulus side sensitivity relative to choice side sensitivity, and low otherwise. A neuron group *α* with 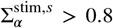 (**Fig.** 2d) is defined as a stimulus processor (1/208 regions, dark blue regions in **Fig.** 2e,f). We plot the distribution of Σ^stim,*s*^ (**Fig.** 2e) of the Beryl regions and observe the stimulus processing regions forming a cluster that have much higher Σ^stim,*s*^ values than the rest of the regions, justifying our categorization. Furthermore, considering that neural signals for the choice variable are relatively more delayed than for stimulus, we can loosen the control for stim-choice correlation during the early stimulus period and only control for stim-prior correlation by using 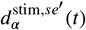 (Eq.3) and 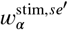 to replace 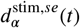 and 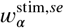 in Eq.4 to calculate a 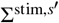. This process results in an expanded list of stimulus response regions (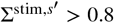 colored in light blue in **Fig.** 2e,f, resulting in 22 additional or 23/208 stimulus side processing areas), and includes visual cortex (VISp, VISam, VISal) and thalamic/hypothalamic (LGd, LP, SGN, ZI) regions that have been previously identified to be relevant for stimulus processing ^1^.

Next, to differentiate integration/choice processing from movement initiation, we define a two-dimensional classification space. We reason that both movement initiation and chocie/integration areas should exhibit significant choice side sensitivity. And definitionally, movement initiators should not be declining in response at movement onset. Thus, we define the quantity Σ^stim,*m*^ ∈ [0, 1]:

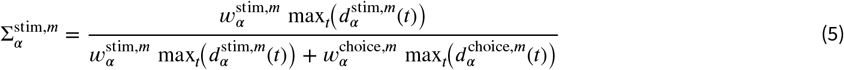

which is small when neurons have low stimulus side sensitivity and high choice side sensitivity, and we define a ‘pre-movement monotonicity’ measure (=1 if the maximum average amplitude 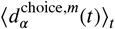 on a sliding 5-bin window occurs in the last five time bins (≈ 10.42 ms) before movement, and the slope > 0.05 in the last twenty bins (≈ 41.67ms); the pre-movement monotonicity is 0 if 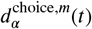 only satisfy the requirement that its slope in the last twenty bins > 0.05, meaning that 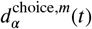 has a ramping shape overall but decays before the last 5 time bins before movement onset).

Combining 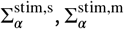 and the pre-movement monotonicity classification (**Fig.** 2d; regional 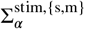 and monotonicity classifications are in **Fig.** S1), we can assign areas to three functional groups: stimulus processors with 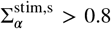 during stimulus period, movement responders with 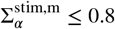 and having monotonicity, and putative integrators/ choice processors (defined separately for stimulus and choice periods) including all the non-movement responders with 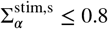 or 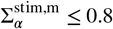, respectively for stimulus and choice periods. The 102/208 Beryl areas with no significant stimulus or choice sensitivity are not assigned a function. The resulting classification of Beryl regions is summarized in **Fig.** 2f.

Plotting the distribution of 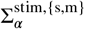 and monotonicity metric values obtained for per-region mean responses in the stimulus and choice periods, we find multi-modal distributions, meaning that there is a clustering in function, **Fig.** 2e: some regions’ responses score as predominantly stimulus-modulated and others as predominantly choice-modulated with a movement ramp. There is a well-separated mode for responses intermediate between stimulus and movement, which we call putative integration/choice areas (or integration areas, for short). Stringent conditioning and significance testing yields a sparse set of stimulus side processing areas (1/208 of recorded areas). We find a less-sparse set of movement initiators (23/208 of recorded areas), and a widespread set of putative integration/choice areas (60/208 of areas), **Fig.** 2f, consistent with previous analyses showing sparse coding of stimulus and widely distributed coding of choice ( ^1^).

Finally, areas differentiating block side during the inter-trial interval are sparse, **Fig.** S1b, S2b. The remaining analysis on prior coding will focus on the time window during trials with the presence of stimulus, to investigate prior effects on sensory, integration, and movement processing.

### Possible mechanisms for prior expectation to influence sensation, integration, and movement

We hypothesize that there are two primary ways the prior input could modulate decision-making: it could inject a differential initial bias between left and right representations, or it could differentially modulate the gain of processing left and right inputs. These effects could be present in stimulus processors, integrators, or movement initiators, **Fig.** 3a, defining six independent mechanisms by which the prior might influence processing during a trial. We examine prior effects by considering how the prior influences the L,R stimulus and choice coding through the following:

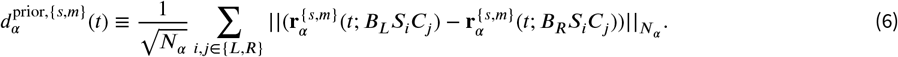

**Figure 3.**
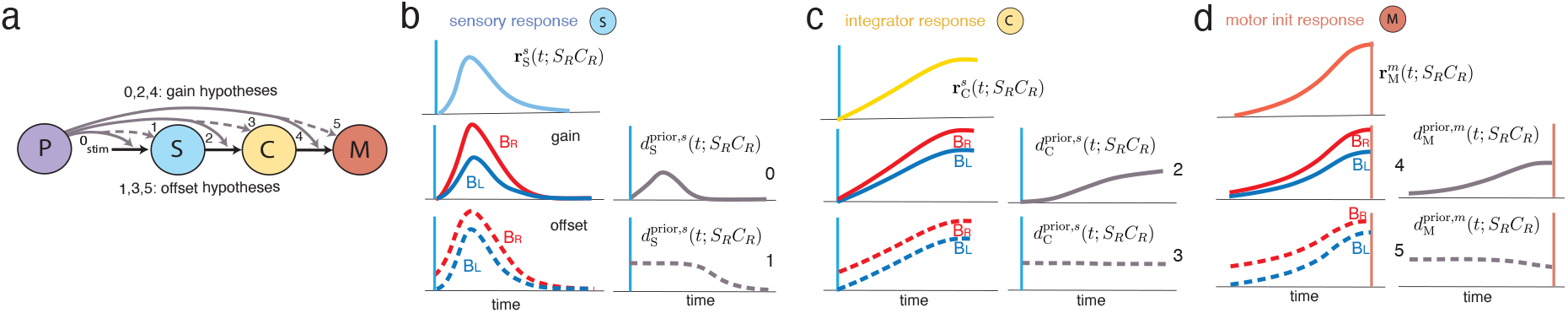
Six hypotheses about expectation-based response modulation and their differential predictions. **a** We summarize six first-order prior modulation hypotheses in this diagram. **b**-**d** Example schematic response curves for sensation, integration/choice, and movement responders of right stimulus and right choice side (left column, top), the responses modulated by the prior, assuming a gain modulation (left column, middle) or bias modulation (left column, bottom). The stimulus, integrator/choice or movement initiation response differences for left versus right prior (right column), for gain and bias modulation (top and bottom, respectively). Note that the difference between gain and bias effects is maximally discernible at the beginning of the response, where the gain modulation is zero and the bias modulation is non-zero.

Conceptually, the encoding of L versus R sensory, integrator/choice, and movement responses, **Fig.** 3b-d (top rows), may be gain modulated by the prior, corresponding to a multiplicative effect on the distance trajectories, **Fig.** 3b-d (middle rows), or the prior might modulate their starting states, corresponding to an initial bias in the direction of the prior, **Fig.** 3b-d (bottom rows). Thus, we consider six independent first-order effects (and any combination of these effects) for how the prior might affect the decision-making process.

### Effects of prior modulation in Beryl regions

The prior or block side *B*_*i*_ could variously be defined by the true block side or different subjective estimators based on the trial history. To maximize the possibility of identification of prior modulation effects in neural data, we choose the definition with strongest representation in the brain-wide responses ^2^, the subjective action kernel. In the stimulus and choice periods, after controlling for correlations by conditioning, we identified 49/208 recorded areas as modulated by prior side during the trial, with a threshold of *p* ≤ 0.01 and FDR correction, **Fig.** 4a (second and fourth rows). The areas differentiating prior side, if using the true block side definition, were qualitatively consistent though much sparser: 24/208 recorded areas with *p* ≤ 0.05 and 8/208 with *p* ≤ 0.01, **Fig.** S3. Curves were assessed for significance by comparison with trial-label shuffle controls, generated by randomly shuffling left, right block assignments for each stimulus, choice condition.

**Figure 4.**
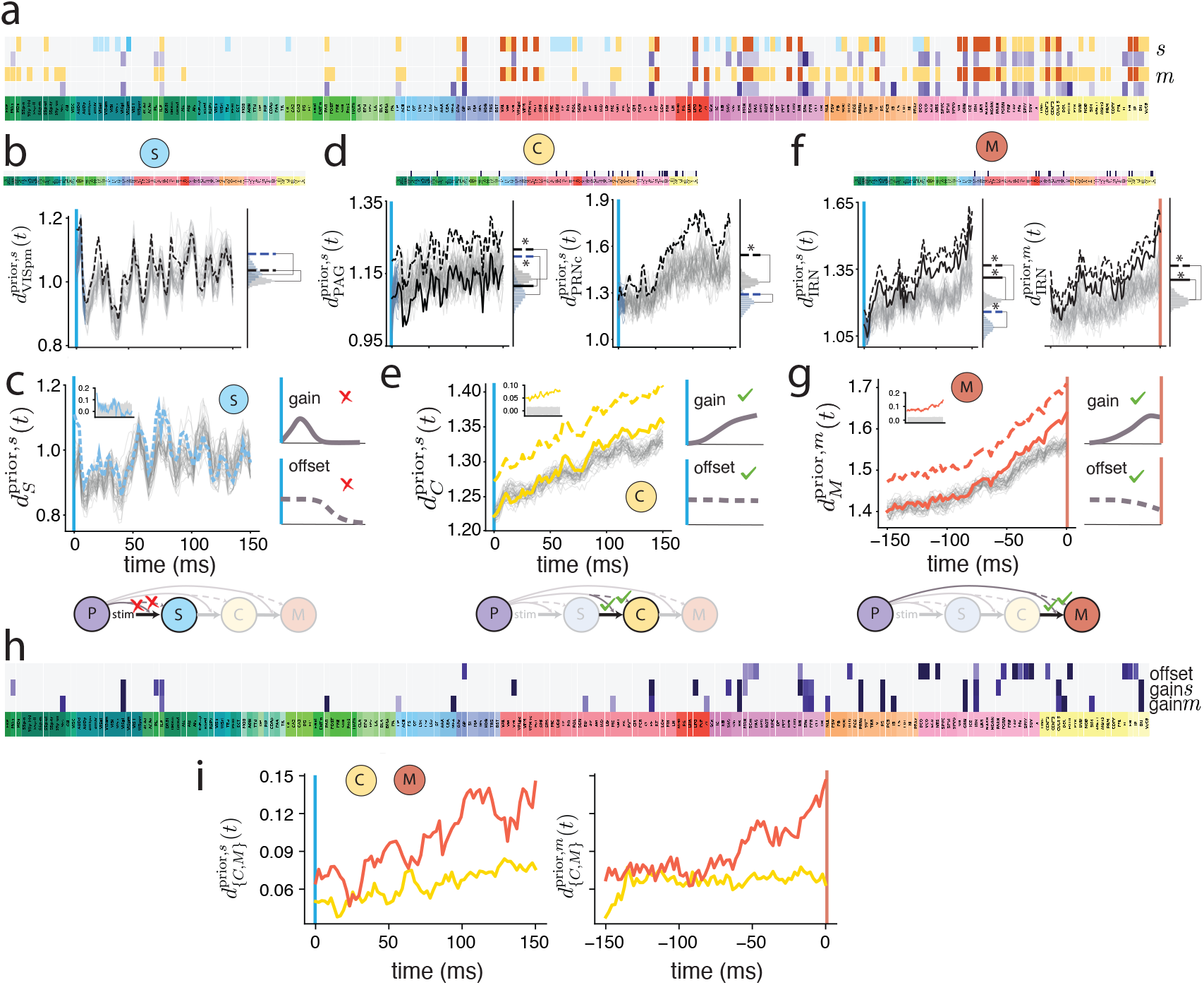
The nature of prior modulation in stimulus processing, integration/choice, and movement initiation areas. **a** Functional classification of regions (blue, yellow, red for stimulus side (S), integration/choice (C), and movement (M), respectively, as in **Fig. 2d-f**) together with prior side sensitivity (purple: regions with significant prior difference, *p* ≤ 0.01 with FDR correction; gray: not significant for prior modulation), aligned to stimulus and movement onset. Lighter blue indicates less stringent inclusion criteria for stimulus side processing (only controlling for prior side but not choice side for early stimulus sensitivity). Darker purple indicates larger effect size 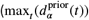 for prior. (**b, d, f**) Top: intersection of S, C, M areas, respectively, with prior side sensitivity (non-zero intersection = black; null = light gray). We use the more stringent definition of S regions (darker blue) in **a**, and results with the lighter blue regions are in **Fig. S4**. Bottom: Prior distance curves (dashed black) for example S, C, M areas, respectively, with shuffle controls (block side labels randomized as in Eq. 8, light gray curves). Solid black curves are the dashed curves subtracting prior bias size, which is defined by the difference between the early-time prior curve mean (first 5 time-bins) and the mean of the early-time shuffles (colored in blue), if a significant bias exists (thus the solid black curve lies behind the dashed curve in **b**). Right panels: Prior distance curve average (black dashed line), distance curve early-time average (blue dashed line; from first 5 time-bins), bias-subtracted distance curve average (black solid line; overlaps with black solid line if bias is not significant), shuffle curve averages (gray histogram), shuffle curve early-time averages (blue histogram). We show stimulus-onset aligned example prior distance curves to examine prior biases (not significant in **b**, significant in 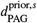 in **d**, 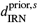 in **f**), and both stimulus aligned and movement aligned prior distance curves to examine gain effects (not significant in **b**, significant for 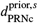 in **d**, 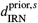 and 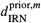 in **f**). p-values: VISpm: overall *p* = 0.0371 (dashed black vs gray hist), bias *p* = 0.029 (dashed blue vs blue hist), gain *p* = 0.992 (solid black vs gray hist). PAG: overall *p* = 0.0058, bias *p* = 0.0005, gain *p* = 0.940; PRNc: overall *p* = 0.0036, bias *p* = 0.097, gain *p* = 0.0005; IRN: overall stimulus aligned *p* = 0.0036, bias *p* = 0.006, gain *p* = 0.0005, overall movement aligned *p* = 0.0037, gain *p* = 0.001. All p-values for individual regions are corrected for false discovery rate (FDR). (**c**,**e**,**g**) Left: Same as (**b**,**d**,**f**), but the colored curves average over all areas with a common function classification (S,C,M, respectively), regardless of individual area significance. S: overall *p* = 0.0315, bias *p* = 0.0025, gain *p* = 1.0; C: overall *p* = 0.0005, bias *p* = 0.0005, gain *p* = 0.001; M: overall *p* = 0.0005, bias *p* = 0.0005, gain *p* = 0.006. Because analysis is by aggregated response there is no FDR correction. Insets: dashed prior distance curves subtracting the means of gray shuffles during each time bin; gray shading indicates the 99th percentile in the shuffles. Right: comparison of the data with schematic curves for hypothesized prior modulation mechanisms. Bottom: mechanisms with statistical support in the data are indicated by checks. **h** Parsing of all regions with significant prior modulation (from **a**) into those showing stimulus-aligned bias modulation (top row), and gain modulation (middle row), and movement-aligned gain modulation (bottom row). Purple: *p* ≤ 0.01 (no FDR correction); gray: not significant. The union of purple areas in the top two rows corresponds to the second row of **a**; the union of the top and bottom rows corresponds to the last row of **a. i** Stimulus and movement-aligned prior distance curves (left and right panels) for integration/choice areas (yellow curves) and movement initiation areas (red curves), superposed for comparison, after bin-by-bin subtraction of the mean of shuffle controls for each curve. The left panel integrator/choice curve is the same as the inset in **e**; the right panel movement curve is the same as the inset of **g**.

We found no regions that exhibit stimulus side processing and prior sensitivity in that response, according to the strict definition (see previous section) of stimulus processing areas (0/1 sensory side processing areas have significant prior sensitivity using the threshold *p* ≤ 0.01; VISpm shows a weak prior sensitivity, *p* = 0.0371), **Fig.** 4a (first two rows) and **Fig.** 4b (top, showing regions with sensitivity overlap), and a small overlap (3/35 areas) by the more permissive definition of sensory processing areas, which upon further examination we find their prior sensitivity does not overlap with early stimulus response (details in SI, **Fig.** S4). By contrast, a broad set of regions exhibits significant integrator/choice processing with prior sensitivity in those responses (14/60 integration/choice areas when aligned to stimulus and 21/60 when aligned to movement onset), **Fig.** 4d (top). Similarly, 14/23 movement initiation areas are prior sensitive, **Fig.** 4f (top). Thus, while correlates of downstream computations for decision making are significantly modulated by the prior, we do not find statistically significant evidence of prior modulation in the stimulus side processing areas on the IBL task.

To quantify the prior modulation effects we measure initial response biases in the sensory and integration/choice areas near the time of stimulus onset. We define the prior bias 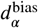 as the average difference between 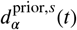 and the mean of shuffle controls in the first five time bins (≈ 10.42 ms) following stimulus onset, **Fig.** 4b,d,f; *p*-values are based on comparison with the distribution of shuffle controls. Differences in response gain induced by the prior, by contrast, would result in small deviations in the difference curves at the start of the trial, but these deviations should increase over time. We therefore assess prior-induced gain modulation in the bias-subtracted curve 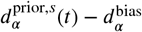 over all time bins after stimulus onset excluding the bias interval. As expected from the lack of prior sensitivity in primary sensory cortical region VISp (**Fig.** S4b (middle)), a key area for sensation, there is no significant bias modulation and no gain modulation from the block prior in sensory response, **Fig.** 4b (bottom, right). The set of sensation regions taken together exhibit no significant bias or gain modulation, **Fig.** 4c.

Integrator/choice regions such as midbrain region PAG and hindbrain region PRNc often exhibit prior sensitivity during the trial, and their responses are significantly modulated by the prior through either bias (for PAG) or gain (PRNc), **Fig.** 4d (bottom). In fact, all integrator/choice areas with prior sensitivity exhibit only gain modulation (e.g. MRN), or are only affected in their initial bias (e.g. PF). The aggregated response of all integrator/choice areas shows large effects in gain and bias, **Fig.** 4e. For movement initiation areas, we consider stimulus-aligned responses to examine initial bias modulation, and movement-aligned responses to examine prior modulation of the gain, **Fig.** 4f. Here also, we find that movement initiation areas have significant prior modulation of the initial bias and the response gain, **Fig.** 4f-g. We also examined the prior L, R trajectories 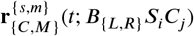 in dimensionally reduced 3-D state-space (**Fig.** S5 b), and they show a clear initial separation that increases over time, consistent with the observation of bias and gain effects on integrator/choice and movement areas. The resulting bias and gain effects across all recorded regions are summarized in **Fig.** 4h. Among the integrator/choice regions whose responses are affected by the prior, 9/24 show bias effects and 15/24 show gain modulation. Among the movement initiation areas modulated by the prior, 8/14 show bias effects while 10/14 show gain effects.

Finally, we compare the magnitude of bias and gain effects in integrator/choice and movement initiation and find that the movement initiation areas exhibit clearly larger bias and gain effects, **Fig.** 4i. We cannot determine from the present analysis whether this difference arises from additional prior-driven modulation of movement areas, or from feedforward amplification of the integrator/choice inputs. Later, we will build a mechanistic circuit model to disentangle the two possibilities.

In sum, the region-wise analyses show notable prior knowledge effects in integrator/choice and movement areas by modulation of both biases and response gains, **Fig.** 4e,g. However, we find no significant evidence of bias or gain modulation in sensation, **Fig.** 4c. We find qualitatively similar results using the true block prior, **Fig.** S3.

### Effects of prior modulation on functionally defined cell groups

Our analyses above were conducted on the level of brain regions. However, results of existing and concurrent work on brain-wide responses in the IBL task show that regional boundaries need not coincide with functional specializations ^1,21^ and individual regions contain neurons with diverse functional response properties. To isolate the effects of prior expectations on functional responses, we partition individual cells into functionally similar types – which are functionally less heterogeneous than anatomical groupings ^21^ – and there examine the role of prior expectations.

To do so, we apply the Rastermap sorting algorithm ^20^ as used in ^21^ to the matrix of per-cell activity feature vectors. For each cell, the activity feature vector is constructed by concatenating together the trial-averaged time-varying responses of a cell for various trial types. Rastermap determines a cellular ordering based on feature vector similarities; the functional clusters are unrelated to region boundaries. This visualization suggests groupings of neurons by function, including integration/choice and movement, **Fig.** 5a (colored boxes highlight some relevant example groupings) ^21^. The stimulus and choice sensitivity of the identified integration/choice (C) and movement (M) groupings are shown in **Fig.** S6, with a nearly linear ramp for the C group culminating in a peak that drops before movement onset, and an accelerating ramp that peaks at movement onset for

**Figure 5.**
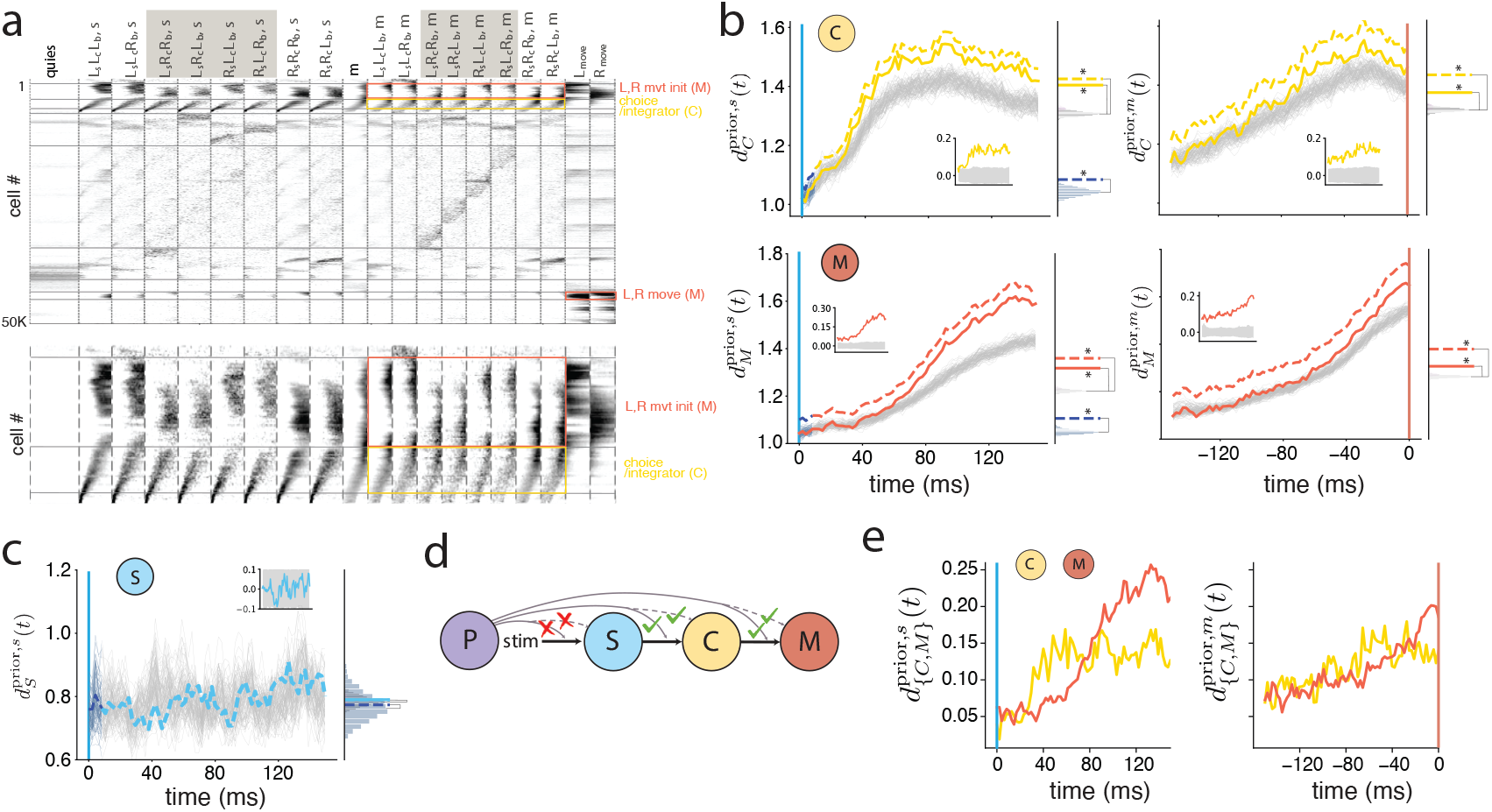
Prior effect on cellularly-defined, region-independent response groups. **a** Top: Individual assignment of the ∼ 60*K* neurons to 2 functionally defined response groups, integration/choice (C) and movement/movement initiation (M) using the Rastermap sorting algorithm ^20,21^ . Bottom: magnification of subset of neurons from above. **b** Prior sensitivity curves for integration/choice (yellow dashed) and movement initiation (red dashed) neuron groups, aligned to stimulus and movement onset (left and right, respectively). Gray curves are shuffle controls. Solid curves are the dashed prior curves subtracting the bias size, which is defined by the difference between the early-time prior curve (first 5 time-bins, dashed blue) and the mean of the early-time shuffle controls. Right panels: distribution of shuffle curve means (gray histogram), distribution of early-time shuffle curve means (blue histogram), and the time-average of the prior distance curve during early-time (dashed blue); time-average of prior distance curve without (yellow or red dashed) and with (yellow or red solid) subtraction of the bias. Insets: dashed prior distance curves subtracting the means of gray shuffles at each time bin; gray shading indicates the 99th percentile in the shuffles. **c** Same as **b**, for stimulus side responding neurons S, blue dashed, defined separately by sorting to find the most stimulus-responding and least choice-responding cells in previously identified sensory regions (Methods and **Fig. S6**). S, C and M groups’ stimulus and choice response curves are shown in **Fig. S6**. Here, because no significant initial bias from prior is identified, the bias-subtracted curve lies behind the dashed curve. p-values (not FDR corrected): C neuron group: overall *p* = 0.0005, bias *p* = 0.0025, gain *p* = 0.0005; M neuron group: overall *p* = 0.0005, bias *p* = 0.0015, gain *p* = 0.0005; S neuron group: overall *p* = 0.3913, bias *p* = 0.4348, gain *p* = 0.4038. **d** Together, the cellular-level results suggest that the prior modulates the bias and gain of C neurons but not S neurons; prior modulation effects are also present in the bias and gain of M neurons. **e** Stimulus and movement-aligned prior distance curves (left and right panels) for C neurons (yellow curves) and M neurons (red curves), superposed for comparison, after bin-by-bin subtraction of the mean of shuffle controls for each curve. The left panel curves are the same as the insets in the left column of **b**, and the right panel the same as the insets in the right column of **b**.

M. Though we do not directly de-correlate responses through trial type conditioning as in the earlier analyses, clustering on response vectors that break out each of the eight trial conditions yields a functional partitioning of cells based on distinctions in their trial type responses. Next, we apply Eq. 6 to these functional groupings (*α* now designates cells within a functional cluster) to determine how the prior affects the responses of C and M neuron groups, **Fig.** 5b. We find that the responses of C and M neurons exhibit strong prior modulation, with significant bias and gain effects, consistent with our findings at the regional level. We also examined the prior L, R trajectories in state-space, **Fig.** S5 a, and they show a clear initial separation that increases over time, consistent with our observations of bias and gain effects on C and M neurons. The regional participation ratio of the top prior-differentiating cells in the C and M neuron groups (summarized in **Fig.** S7, S8) shows wide-spread prior signals across the whole brain, with leading regions from the hindbrain, striatum, thalamus, and hippocampus for C neurons, and the hindbrain, cerebellum, and somatosensory cortex for M neurons.

Next, we consider how sensation-related neurons are modulated by the prior. The Rastermap-based approach identifies a cluster of neurons with short-latency responses to trial onset and feedback onset events, the former of which involves visual and auditory stimuli, while the latter involves an auditory (and in rewarded trials, a slower gustatory) stimulus. On examination, these neurons have a sub−40 ms latency that is inconsistent with typical visual response latencies and do not exhibit stimulus side sensitivity, suggesting they are responding to the auditory click at each event. Instead, we sorted all neurons in areas known to be relevant for visual processing ^1^ based on their stimulus and choice responses, and identified a group of cells with the most different response for stimulus and the least different response for choice. These neurons exhibit sensory response transients with ∼ 40 ms response latencies and stimulus side sensitivity, **Fig.** S6. Applying Eq. 6 to these sensation neurons, we find no significant prior modulation in either their response bias or gain (**Fig.** 5c), again consistent with our findings at the regional level.

In sum, the regionally-independent cellular analysis shows strong bias and gain modulation of both integration/choice and movement neurons by prior expectations, and little evidence of prior expectations modulating the biases or gains of sensation, **Fig.** 5d. These region-independent results are consistent with the analyses performed at the regional level. Also consistent with the regional analysis, when we compare the magnitude of prior effects in the integration/choice and the movement functional groups, we find that the movement initiation group exhibits larger bias and gain effects compared to the integration/choice group (**Fig.** 5e). The consistency of region-wise and cell-wise results provides a measure of confidence in the robustness of conclusions about the loci and types of modulation exerted by prior expectations. Next we build a mechanistic circuit model of the sensorimotor arc of decision making and possible locations, times, and ways in which the prior could modulate these dynamics to attempt to understand which mechanisms are present in the brain.

### Emergent simplicity: A mechanistic circuit model

Despite the possibility that recording increasingly many neurons across the brain during a task would make the picture of how activity underpins behavior increasingly complex, and indeed recording more neurons reveals functional categories of activity that were previously unanticipated ^21^, we also find that there is an emergent simplicity. We will show below that it is possible to reproduce task-relevant brain-wide neural responses and resulting behavior with a simple mechanistic neural circuit model consisting of 8 nodes. Previous mechanistic modeling of how the brain might solve the IBL task involved training a recurrent neural network on the task ^22^, and reducing the trained network into a two-neuron effective circuit representing within-trial stimulus integration/choice and slower across-trial prior estimation; however, though it reproduced animal behavior it was not compared to neural data ^22^. Here we ask whether, if we instead begin with a simple mechanistic model, it will succeed in fitting large-scale neural activity, and whether that neural activity-fit model would then also reasonably capture behavior. If it were possible to construct such a model, it would yield mechanistic insight about the brain’s task-related computations and also allow us to estimate more quantitatively the effects of prior modulation described above.

Our simple circuit model is a bilateral version of the distilled two-neuron integrator-prior circuit from ^22^, with the addition of a pair of sensory processing units and movement initiation units, as well as a final action readout. Each of the eight units, to be understood as coarse-grained representations of a group of neurons participating in the corresponding processing, excites itself and inhibits the opposite unit, **Fig.** 6a, and is coupled to prior units as specified in the following:

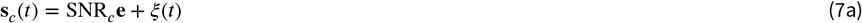

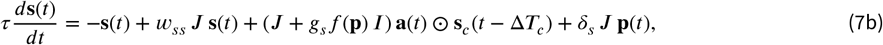

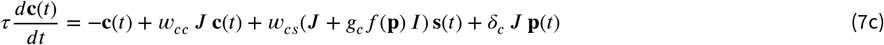

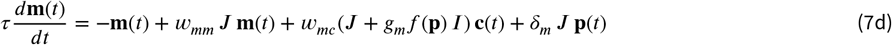

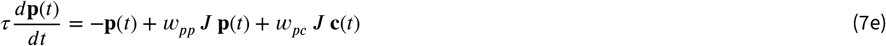

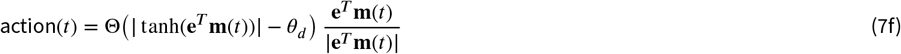

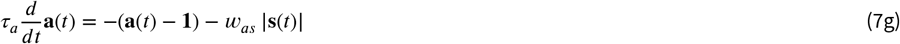

where **s**_*c*_(*t*) is the noisy sensory drive received by the sensory processing units **s**, Eq. 7a, the ±1 entries of **e** designate a left or right input, the parameter SNR_*c*_ specifies amplitude as a monotonic function of stimulus contrast (c), *ξ* is a unit-norm zero-mean Gaussian, ⊙ is element-wise multiplication,*I* is the identity matrix, **e**^*T*^ = (+1 −1) and 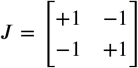. The sensory processing units **s**^*T*^ = (*s*_*L*_, *s*_*R*_), Eq. 7b, respond to noisy stimuli **s**_*c*_(*t*) with contrast-dependent temporal delays Δ*T*_*c*_. The sensory units therefore represent the perceived stimulus; they adapt through coupling with adaptation units **a**^*T*^ = (*a*_*L*_, *a*_*R*_)(Eq. 7b,g) to create short-lived stimulus responses. The integrator/choice processing units **c**^*T*^ = (*c*_*L*_, *c*_*R*_), Eq. 7c, receive inputs from the stimulus processing units with weight *w*_*cs*_. They integrate their inputs over time through recurrent self-excitation *w*_*cc*_ ^23^, and send their outputs to the movement and prior processing units with weights *w*_*pc*_ and *w*_*mc*_, respectively. The movement initiation units, **m**^*T*^ = (*m*_*L*_, *m*_*R*_), Eq. 7d, and prior units **p**^*T*^ = (*p*_*L*_, *p*_*R*_), Eq. 7e, self-excite with weights *w*_*pp*_ and *w*_*mm*_ respectively. The prior units represent the estimated block side and can inject a bias into all the other units through the inputs *δ*_*i*_ > 0. They can also modulate the weights between sensory, integrator/choice and movement units through the gains *g*_*i*_ > 0 multiplied by *f* (**p**) = sgn(**s**^*T*^ *J* **p**) | **e**^*T*^ **p**|, which has a magnitude |*p*_*L*_ − *p*_*R*_| and is positive (negative) if the trial is con-(dis-)cordant, defined by whether the stimulus **s** and prior **p** are on the same (opposite) sides; here, the prior effect is to boost the gain of stimulus inputs to the sensation, integration/choice, and movement nodes in concordant trials, and suppress it on discordant ones. Finally, when tanh(|*m*_*L*_ − *m*_*R*_|) > *θ*_*d*_, an action is taken in the direction specified by that difference, Eq. 7f.

**Figure 6.**
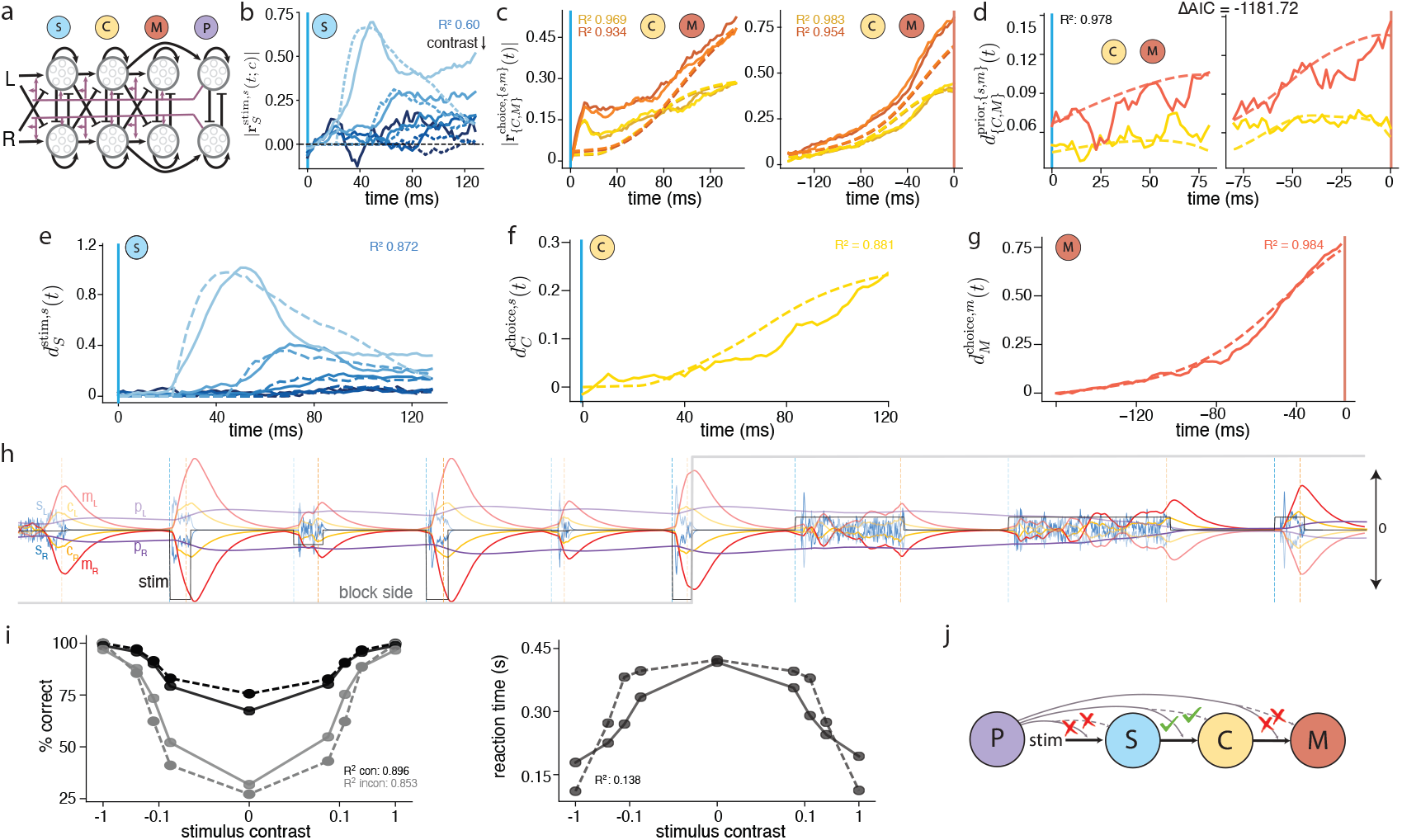
Simple mechanistic circuit model of brainwide dynamics on the IBL task. **a** The model consists of a bilateral pair of units for stimulus (S), integration/choice (C), movement (M), and prior (P) processing. Units excite ipsilaterally and inhibit contralaterally. S unit responses drive the C units, which drive the M and P units. P units inject biases into all other units and modulate the weights between S, C and M units. Each unit is a coarse-grained representation of a group of neurons participating in the corresponding function. **b-d** Finding model parameters by fitting simulated model outputs to aggregated root mean square response amplitudes of S, C, and M areas (**b-c**), and the aggregated prior sensitivity curves of C and M areas (**d**) (from regional analysis). We report ΔAIC = −1181.72 as compared to the baseline model without any prior modulations in **Fig. S9a**. Dashed lines: model; solid lines: data. Fits for S are to all five contrasts, C and M for left/ right choices. Sensory responses (**b**) are aligned to stimulus onset and data are shifted by subtracting baseline activity (average of the first 7 time bins) to align with the model. C and M are aligned to stimulus onset (left panel in **c-d**) and movement onset (right panel in **c-d**) times, and data are shifted to align with the model at stimulus onset. Prior sensitivity data curves are the same as in **Fig. 4i** but truncated to 80 ms aligned to stimulus/movement onset. **e-g** Comparison of the fitted model’s stimulus side sensitivity (L-R euclidean distance), choice side sensitivity, and movement sensitivity predictions, respectively (dashed lines), with data (solid lines). The model prediction curves are each multiplied by a global scaling factor on the order of 1 to match the data. **h** Prediction of temporal response trajectories for S (pale and darker blue curves, for left and right, respectively), C (pale and darker yellow), M (pale and darker red), and P (pale and darker purple; values multiplied by 10 for visualization here) from the fitted model’s units over several trials, black lines indicating signed stimulus contrast and gray lines the block side. Stimulus onset (vertical blue dashed line) ultimately results in movement onset by the model (vertical orange dashed line), followed by a short fixed-duration action period (40 ms) and a long (1 s) fixed-duration inter-trial period (combined duration: orange to next blue dashed line). Note that any prior biases into S,C, or M are equal and opposite for each contralateral pair, thus those curves remain symmetric about 0. **i** Behavioral predictions by the fitted model (dashed lines) match animal behavior (solid lines) (model behavior prediction is obtained from fitting to physiological responses in **b-d** with no fitting of the model to behavioral data). Left: percentage of correct choices in congruent (same trial and prior sides) and incongruent (opposite trial and prior sides) trials, as a function of log stimulus contrast. Right: reaction time (interval from stimulus onset to movement onset) as a function of log stimulus contrast. **j** Combined conclusion from data and mechanistic model: prior expectations directly modulate integration/choice response biases and gains. There is no statistically significant evidence of direct prior modulation of stimulus areas, and strong modulation effects in movement area responses are explained in the model by amplification of inherited integration/choice inputs, thus we find no evidence of additional direct prior modulation of movement initiation areas.

We simulated Eq. 7 with the same statistics of inter-trial intervals, block structure, and stimulus sides as in the experiment (details in Methods) using a neural time constant *τ* = 40ms. We found values for all weight, gain, bias, and action threshold parameters by fitting the model to the aggregated root mean square response amplitudes of sensory, integrator/choice, and movement-initiation areas, and the aggregated prior sensitivity curves of choice/integrator and movement areas (from regional analysis), **Fig.** 6b-d (full set of fitted parameter values listed in Methods). We did so in two stages: first, fitting contrast-dependent stimulus responses to extract relevant parameters (SNR and delay parameters, adaptation time constant *τ*_*a*_ and stimulus weights *w*_*ss*_, *w*_*as*_), which achieves *R*^2^ = 0.872, **Fig.** 6b); next, obtaining the remaining parameters (other weights, action threshold *θ*_*d*_, and the prior modulation parameters *δ*_*i*_, *g*_*i*_) by fitting the model’s choice-related responses and prior distance curves, with the integration weight *w*_*pp*_ constrained to a narrow range to ensure proper long-term prior integration. Parameters *g*_*s*_ and *δ*_*s*_ were set to zero based on the absence of significant prior-related differences in stimulus-response regions (**Fig.** 4c). The model achieved strikingly good fits, with 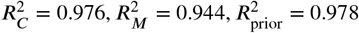, **Fig.** 6c-d (fitting procedure details in Methods).

These fits yielded the following feedback and prior modulation strengths: *w*_*ss*_ = 0.00, *w*_*cc*_ = 0.43, *w*_*mm*_ = 0.27, *w*_*pp*_ = 0.496, which generate interpretable values of integration time-constants at each stage without having explicitly fit those parameters, corresponding to a sensory processing time-constant of *τ*/(1 − 2*w*_*ss*_) ≈ 40ms, an integration/choice integration time-constant of *τ*/(1 − 2*w*_*cc*_) ≈ 270ms, a movement-initiation ramping time-constant of *τ*/(1 − 2*w*_*mm*_) ≈ 90ms, and a prior integration time-constant of *τ*/(1 − 2*w*_*pp*_) ≈ 5s. These obtained time-constants are consistent with rapid sensory responsiveness, the role of the integrator/choice processors in estimating the stimulus side through integration over the ≈ 0.2 − 1 s duration of the trials, and with the previously found prior integration kernels of ≈ 10 trials and ≈ 5 − 10s ^1,2^. The stimulus processing and movement initiator time-constants by contrast are substantially faster, as one might expect. Additionally, the fit parameters for input gain into the integration/choice processors and movement initiators are *w*_*cs*_ = 0.17, *w*_*mc*_ = 0.50, corresponding to an overall input amplification of *w*_*cs*_/(1 − 2*w*_*cc*_) ≈ 1.2 by integration/choice processors and *w*_*mc*_/(1 − 2*w*_*mm*_) ≈ 1.1 by movement initiators.

We can use the model to interpret the strengths and types of influence that the prior exerts on processing at different stages of the IBL task. In the model above, *g*_*c*_ ⟨|*f* (**p**) |⟩ = 0.197, corresponding to a ≈ 20% modulation in the gain with which integration/choice processors integrate their sensory inputs. By contrast, we find that *g*_*m*_ ⟨|*f* (**p**) |⟩ = 1.14 × 10^−3^, corresponding to a ≈ 0.1% or negligible modulation in the gain with which movement initiators integrate their inputs. The prior modulated the initial bias of integration/choice processing by the fractional amount *δ*_*c*_ ⟨|**p**|⟩ /(*w*_*cs*_ range( **s** )) = 0.108, or ≈ 11% of the total inputs. The prior injects *δ*_*m*_ ⟨|**p**|⟩ /(*w*_*mc*_ range(|**c**|)) = 1.267 × 10^−6^ or ≈ 1.2 × 10^−4^% of the total inputs driving movement initiation, again a negligible quantity. In sum, both data and model show pronounced prior effects on both gain and bias in integration/choice processing. And, though both data and model curves appear to show even more pronounced effects of the prior on movement initiation (**Fig.** 4i), the model indicates that these effects are inherited by the movement processors from their integration/choice inputs and the greater effect size is consistent with the overall input amplification by the movement processors, rather than with a direct and additional prior effect on either the gain or bias of the movement processors. If we consider a constrained version of the model in which we remove prior gain and bias modulation terms for the movement initiator and refit all parameters, the result is a comparably good fit to the physiological data as the full model (fits for curves as in **Fig.** 6c-d yield 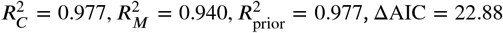 compared to the full model; AIC is the Akaike Information Criterion), supporting the lack of bias and gain modulation of movement initiation in the brain (**Fig.** 6j).

Next, to probe the overall contribution of prior expectation to neural responses and the relative importance of gain versus bias modulation, we re-fit the model with some prior modulation parameters set to zero. With no prior modulation anywhere, fits to the data strongly degrade **Fig.** S9a), as expected from the observed prior effects in **Fig.** 4. Removal of either the prior-induced biases 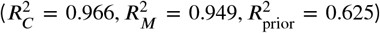 or of the prior-based gain modulations 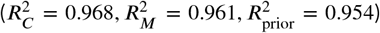 yields worse fits, with a somewhat larger effect from the biases (ΔAIC = 919.26 comparing removal of bias versus gain, **Fig.** S9), possibly because the bias influences the entire duration of the trial while the gain effects are expressed with signal integration latencies of response in integration/choice and even longer latencies in movement areas.

We next test the previously fitted model by considering how it reproduces the data distance curves 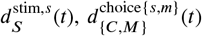. We find that it matches the data curves (up to a scaling factor of order 1 for each curve: 1 for 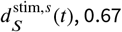 for 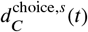, and 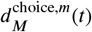, **Fig.** 6e-g (*R*^2^ = 0.872, 0.881, 0.984 respectively).

Further, the fitted model can be used to simulate trial-by-trial neural dynamics, obtaining time-varying stimulus processing, integrator/choice, movement, and prior responses with plausible profiles, **Fig.** 6h; this process also yields behavioral responses. We use these model outputs to quantify its psychophysics, plotting the percentage of correct responses and the reaction time as a function of stimulus contrast. Remarkably, though the few parameters of the model were fit only to neural data, we find that with no additional parameter tuning, fine-tuning, fitting or scaling, the model accurately predicts mouse behavior, **Fig.** 6i. The model reproduces the performance on congruent and incongruent trials (defined as same or different stimulus side from prior side), *R*^2^ = 0.896, 0.853, including the performance gap on low-contrast trials where prior information is critical for performance. It also predicts the reaction time, defined as the time between stimulus onset and movement onset within a trial as a function of contrast, *R*^2^ = 0.138.

Finally, we can further test the model’s conclusions about prior modulation. Above we did so based on its match to physiological data, now we also do so by comparing the model’s psychometric and chronometric behavioral predictions with animal behavior (as above, we fit the model only to physiology not to behavior, using the behavior only as a final test). With no prior modulation, the model’s concordant and discordant trial responses are identical to each other, poorly predicting mouse behavior on low-contrast trials (**Fig.** S9a). The model better predicts behavior with both bias and gain modulation than with only one of them (**Fig.** S9b-c). The best behavioral predictions are obtained when the gains and biases of integration/choice units are modulated by the prior, **Fig.** 6, and further modulation of the movement initiation units does not yield better predictions of behavior. This final set of behavioral prediction results, consistent with the previous conclusions from physiology, provides further support for our conclusion that prior effects are important, that they primarily affect integration/choice responses, and that they do so both by biasing their states and by modulating the gains by which they integrate their inputs.

We briefly mention here a model variant, in which the bias effects (the *δ* terms in the equations above) are replaced by a constant offset of the integration/choice and movement states (alternative model equations and parameters in Methods, Eq.9, results in **Fig.** S10). This alternative model also produces good-quality fits to the physiological data (**Fig.** S10e, 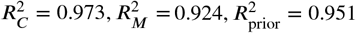) and good predictions of behavior (percentage correct with *R*^2^ = 0.972, 0.976 for congruent and incongruent trials, reaction time with *R*^2^ = 0.452). It yields qualitatively consistent conclusions about the influence of the prior as above, finding a significant influence of the prior on integrator/choice processing in the form of both gain and bias modulation, a lack of significant influence of the prior on stimulus processing, and a lack of additional direct modulation of movement initiation beyond inherited effects from the integration/choice processors (*g* _*c*_⟨|*f* (**p**) |⟩ = 0.149 or ≈ 15% integrator/choice gain effect, *g*_*m*_⟨| *f* (**p**)|⟩= 0.001 or negligible ≈ 0.1% movement gain effect; *δ*_*c*_⟨| **p**|⟩ /(*w* _*cs*_ range( |**s**| )) = 0.307 or ≈ 31% integration/choice bias relative to total input, and *δ*_*m*_ ⟨|**p**|⟩ /(*w*_*mc*_ range(|**c**|)) = 1.078 × 10^−6^ or negligible ≈ 1.1 × 10^−4^% movement bias relative to total input. Also as before, the alternative fit to the physiological data best predicts the psychometric and chronometric curves from mouse behavior when integration/choice processing is subject to both bias and gain modulation by the prior, and when stimulus processing and movement initiation are not (**Fig.** S10d, ΔAIC = −3.86 compared to the full model in S10e). This finding verifies the robustness of our conclusions to model variations.

In sum, we find that a simple mechanistic model of IBL decision-making provides good fits to neural population responses, results in robust and highly interpretable parameters that quantify the loci, types, and amplitudes of prior effects on task processing, and predicts animal behavior (psychometric and chronometric responses) though fit only to physiology. By generating a good match to the difference curves for stimulus, integration/choice, and prior, and a direct prediction of behavior, it elucidates the mechanistic links between these quantities. The model also allowed us to conclude that the primary effect of prior expectations in the IBL task is to modulate the bias and gain of integration/choice processing, and that the large modulation of movement responses by the prior are inherited from integration/choice input modulation rather than an additional direct modulation.

## Discussion

Leveraging the International Brain Laboratory’s unprecedented brain-wide recordings across 208 regions and 60,000 neurons, we examined where, when, and how learned expectations shape ongoing computations across the arc of perceptual decision-making. By disentangling the correlated variables and neural representations of stimulus, choice, and prior knowledge, we found that rather than reformatting early sensory codes, prior knowledge biases downstream computation in the IBL task. These findings were robust whether we assigned functional roles to brain regions or neurons, and they support hierarchical-inference accounts in which beliefs and uncertainty modulate decision variables more than early feature encodings ^9,24–29^.

Next, we discovered an emergent simplicity: a compact mechanistic neural circuit with few parameters captured the task-related brain-wide neural dynamics across conditions. With zero additional fitting, this neural circuit activity model then predicted and matched mouse behavior (psychometric and chronometric curves). The model also generated interpretable values of integration time-constants at each stage (sensory processing, integration/choice processing, movement initiation, and the integration of evidence for the prior) without having directly fit those parameters. It provided a critical mechanistic insight that we could not obtain from the data without the model: that the large observed bias and gain modulations in movement initiators are not directly or additionally modulated by prior expectation, but rather are an inherited amplification of the inputs received from the modulated choice/integrator circuits. Thus, the model captures the sensorimotor arc of the decision process, bridging from neural activity to behavior.

We further found that in the IBL task, prior knowledge also does not directly bias movement initiation and movement circuits beyond modulation of their integration/choice inputs. In the IBL task, choice and movement must take into account not only prior expectation but also the current sensory evidence. Thus, the brain must be prepared to take either action based on the evidence even for the same prior expectation, possibly making direct modulation of movement areas by prior expectation sub-optimal. By contrast, in tasks where prior expectations fully predict the action, direct modulation of movement areas by the prior may promote faster reactions without loss of accuracy ^30,31^.

Our findings about the loci and stages at which prior expectations modulate decisions are actionable. They predict specific entry points for learned expectations into the brain’s decision making process that can be tested by perturbation studies. And, the combined standardization of the IBL task and our analysis method generate replicable biomarkers for the study of neuropsychological disorders — e.g., gain modulation ratios and initial-bias indices — that could be compared across species and cohorts including in disease models ^32,33^. Indeed, the present work connects to computational-psychiatry views that symptoms may reflect mis-weighted priors: psychosis has been interpreted as overly-strong or imprecisely targeted high-level priors, while autism has been argued to involve attenuated priors ^34–38^. Our mapping suggests candidate loci of such mis-weightings (in the integration/choice stages), and alternatively allows future investigation of the possibility that psychosis might involve mis-timed or mis-targeted prior modulation at different stages or loci than in neurotypical individuals.

### Relationship to existing work

Our modeling approach is related to the drift diffusion model (DDM) ^39^, as we essentially employ a leaky integrator version of DDM, but with four coupled integrators operating at different time scales (stimulus, integrator/choice, movement initiation, and prior expectation). In addition, we apply the model at the level of neural rather than behavioral dynamics, fitting it to neural population activity and obtaining a good prediction of behavior without any further fitting.

Our result that prior expectations influence decision making through gain and bias modulation aligns with previous behavior modeling work using DDM frameworks ^5,40^ and with analyses of rodent and primate neural data ^7,17–19^. Gain modulation has been observed when previous actions (interpreted as a subjective prior) alter the effective weighting of incoming evidence in humans and mice, and in primate studies showing expectation-or attention-driven increases in population response gain during perceptual decision making ^10,11,13–15^. Complementing this, a parallel line of work demonstrates bias modulation: studies in parietal cortex of monkeys and rodents show that prior or contextual information biases neural responses during perceptual tasks ^17,19^, and human EEG and fMRI studies also report baseline shifts associated with prior expectations that modulate stimulus-evoked activity in perceptual decision-making tasks ^18,40^.

The evidence on whether prior expectations modulate early sensory regions has been mixed ^2,6,8,9,11,13,14^. Our study found no prior modulation of differential sensory responses, while other studies in humans and non-humans report top-down modulation of sensory processing such as the effects of depth and perspective on the perception of object size, and even on V1 responses ^41^. A possible resolution of discrepancies may be that prior modulations are measured at different phases: for instance, the reported V1 modulations in ^41^ appear 150–200 ms post-stimulus rather than immediately, suggesting a delayed re-entrant or top-down influence on V1, likely from downstream areas V3A/V7 which primarily process depth and perspective cues to estimate object size. The lack of immediate modulation of low-level sensory responses is consistent with our findings. Most directly comparable with our analysis, a decoding study of IBL neural activity reported that prior expectation may modulate perception by showing that early visual areas appear to represent the prior ^2^. The likely source of this discrepancy is a difference in controlling for correlated variables. While ^2^ assessed the statistical significance of the decodablity of the prior side from different brain regions, prior expectation is correlated with current stimulus, previous choice, and current choice. Thus, decodability provides an upper bound on the existence and strength of prior effects. We attempted to stringently control for correlated variable effects by analyzing response differences within each single stimulus and choice side trial type. Additionally, we found that many neurons and regions that respond to stimuli at early times also integrate this information to later represent choice. We classified these as integration/choice responses. Many of these regions do show significant prior signals. Our stringent criterion for pure sensory neurons and regions resulted in a small set of sensory neurons and regions; though we found a slightly expanded but still sparse set of pure stimulus response regions when only controlling for prior side during early stimulus sensitivity (VISp, VISam, VISal, LGd, LP, PO,**Fig.** 2f), none of these regions showed significant prior sensitivity during the stimulus period.

### Future Directions

On the microscopic side, future work will be needed to elucidate how the observed macroscopic prior gain and bias modulation effects are implemented. Possible mechanisms include neuromodulatory signals or shunting inhibitory effects for gain change.

On the macroscopic side, it remains to determine causal relationships among brain regions to determine which regions compute and broadcast the prior-based modulatory signals we have identified as affecting computation. This requires perturbation studies, as noted above. It will be especially interesting to investigate how the representation of prior information and its effects on integration/choice processing changes over learning; our analysis methods can be used to quantitatively track the changing influence of the prior over learning, and to quantify the diversity of learning trajectories and dynamics among individual animals. Collectively, our results define the neural circuitry and mechanisms by which prior knowledge is applied to decision-making computations, shifting the understanding of expectation from a high-level influence to a precisely localized and dynamically characterized modulation. The novel synthesis between large-scale population electrophysiology and mechanistic modeling reveals a new organizing simplicity behind cognition across the brain.

## Methods

### Neural Data Analysis

We analyzed neural data using the population trajectory analysis method described in previous work ^1^, applying it to quantify time-resolved differences in brain-region responses under left versus right conditions of stimulus, choice, or block prior. We used the same inclusion criteria as previous work ^1^, including session and insertion quality control, trial quality control (excluding trials with missing events, or with duration shorter than 0.8s or longer than 2s). We also exclude the trials without biased block sides at the beginning of each session. For the regional analysis, a region needs to have at least 20 cells to be included.

#### Significance Testing

We explicitly controlled for correlations among stimulus, choice, and block (prior) variables to obtain a time-resolved difference curve for a given population, 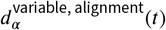, as described in Results. To create a null distribution, we generate 2000 distance trajectories with randomly shuffled L-R labels within the trial type dichotomies (**Fig.** 2a-c semi-transparent gray curves). For example, for stimulus sensitivity, a control distance curve is generated by

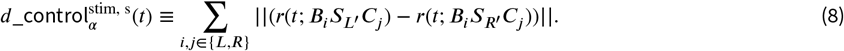

where *L*^′^ and *R*^′^ are shuffled stimulus left and right labels for trials of block side *i* and choice side *j*. We compare the time-averaging value of the measured euclidean distance curves, 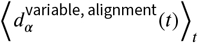 with the time-averaging values of the shuffled control curves, 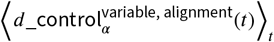 to generate a p-value, 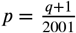, where q is the number of times that the control curves have 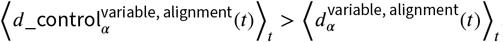. For analysis based on Beryl regions only, we applied the Benjamini-Hochberg method for false discovery rate (FDR) correction over the p-values of all regions included, using the package statsmodels.stats.multitest.multipletests in python. We then used a threshold of *p* ≤ 0.01 to obtain an overall significance score 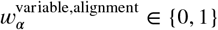 denoting statistical significance for variable x (stim, choice, or prior) in the population.

#### Identifying Bias and Gain Modulation by the Prior

For populations showing significant prior sensitivity during trials, we need to assess whether this effect arises from changes in bias or gain, thereby clarifying the mechanism through which the prior influences processing. To evaluate bias, we focused on the stimulus-aligned time window and compute the time-averaged prior difference curve over the first five bins (≈ 10.42 ms), comparing this value against the corresponding shuffle controls. We define a p-value for bias as 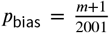, where m is the number of times that the control curves have 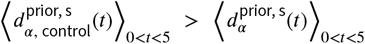. Populations that are statistically significant for the prior during stimulus period and have *p*_bias_ ≤ 0.01 are identified as having bias modulation. We then define bias as the average difference between 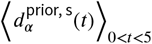 and that of the shuffle controls, namely 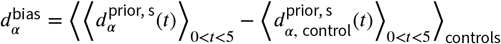 for populations with significant prior biases, and 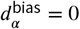 for populations without. To examine if a population is modulated by the prior through gain effects, we subtract the bias from the prior distance curve to get 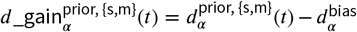, and compare its time average with shuffle controls after the initial bias interval (first 5 time bins) to define a p-value: 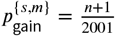, where n is the number of times that the control curves have 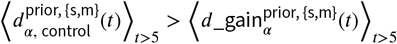. Populations that are statistically significant for the prior and have 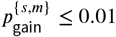 are identified as having significant gain modulation by the prior.

#### Region-independent Analysis

We calculate per-cell activity feature vectors, containing stimulus onset-aligned and movement-aligned trial averaged PETHs for the eight *S*_*i*_*C*_*j*_*B*_*k*_ trial type combinations, as well as pre-stimulus quiescence and post-movement onset PETHs, using the same procedures as in ^21^. Applying Rastermap ^20^ sorting algorithm on these feature vectors generated clearly functionally distinct groups, from which we selected the integrator/choice, movement initiation, and movement generation cells. For the current analysis we do not differentiate between movement initiation and generation, therefore we combine these two categories as the movement cells to investigate prior modulation on them. Both the integrator/choice and movement groups show significant stimulus and choice coding when aligned to stimulus and movement onset (S6). The Rastermap algorithm does not clearly identify a stimulus-response group that only responds significantly to stimulus at early times but not to integrator/choice. We focus the search for stimulus processors in eight selected areas relevant for visual processing, as identified from our analysis and previous work ^1^: VISpm, VISam, FRP, VISp, VISli, LGd, LP, and NOT. We sorted the neurons from these regions by the stimulus differences in their average PETHs ( ⟨*r*^*s*^(*t*; *S*_*L*_*C*_*j*_*B*_*i*_) ⟩ _*t*_ − ⟨*r*^*s*^(*t*; *S*_*R*_*C*_*j*_*B*_*i*_) ⟩_*t*_) between the 20 to 90 ms after stimulus onset, and identified the top stimulus responding cells by taking the union of the top 400 cells from all four *S*_*L*_*C*_*j*_*B*_*i*_,*S*_*R*_*C*_*j*_*B*_*i*_ pairs. We then compute their choice differences ( ⟨*r*^*s*^(*t*; *S*_*j*_*C*_*L*_*B*_*i*_) ⟩ _*t*_ − ⟨*r*^*s*^(*t*; *S*_*j*_*C*_*R*_*B*_*i*_) ⟩ _*t*_) during the same time window, and removed the cells with choice difference larger than 0.5 for any of the four choice pairs. The resulting stimulus-response group contain 79 cells that have significant stimulus coding but not for integration/choice (**Fig.** S6).

### Modeling and Data Fitting

To implement simulations, we discretized Eq.7 with *dt* = 2ms. The maximum duration of a trial is 2s = 1000 *dt*, with an inter-trial period of 1s = 500 *dt*. Trial statistics were matched to those used in the experiments. Contrast levels of 100%, 25%, 12.5%, 6.25%, and 0% were presented with probabilities in the ratio 2:2:2:2:1, respectively. The trials are organized into blocks, where one of the left and right sides is designated as the block side with a biased 80% probability of being the correct trial side, and the opposite side with 20% probability. The block lengths were sampled from a geometric distribution with mean of 60, minimum length of 20 and maximum length of 100 trials. Unlike the experiments, simulations began directly with biased blocks using an 80/20 side probability, without an initial unbiased block. The initial block side was chosen randomly, and subsequent block sides alternate after each block. Each simulation session consists of at least 4 blocks of variable lengths, allowing the dynamics to run continuously for long enough before trial-averaging them to compare with data.

To examine the robustness of our conclusions to what form the prior bias might take, we implemented two different forms of bias: one as in Eq.7, where the bias is an input injected from the prior nodes to the sensory, choice/integration, and movement nodes, and is being integrated along with the actual stimulus input. A second type of bias directly shifts the activities in the sensory, choice/integration, and movement nodes:

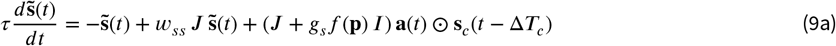

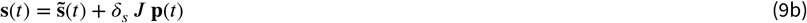

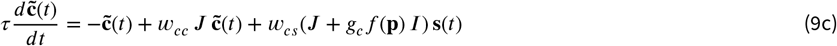

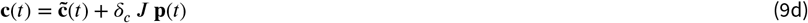

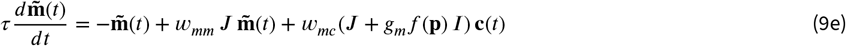

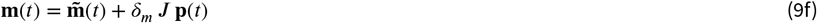

The bias is only activated at 100 ms before stimulus onset until the end of a trial in both cases, so that neuronal activities can decay back to the original zero baseline in the intertrial interval.

We used differential evolution (DE) algorithm for a global search in the logarithmic parameter space, and covariance matrix adaptation evolution strategy (CMA-ES) for local refinement to optimize model parameters. The first part of the fitting process targets stimulus-processing parameters: specifically, the adaptation time constant (*τ*_*a*_), the recurrent stimulus weight and feedforward weight from stimulus to adaptation unit (*w*_*ss*_ and *w*_*as*_), and the parameters governing the functions (of stimulus contrast *c*) for the stimulus signal-to-noise ratio, *α, β* for 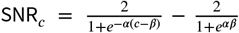, and stimulus response delay, *σ, λ* for Δ*T*_*c*_ = round(*σ*/(1 + *λ c*)) (rounded to integer numbers for discrete simulation). Empirical data consisted of the root mean square of PETHs for trials of each contrast 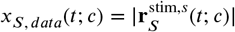, where |.| is the Euclidean norm) from stimulus-responsive regions (VISpm, VISam, FRP, VISp, VISli, LGd, LP, NOT, a set combining results in **Fig.** 2 and previous work^1^) during the first 100 ms after stimulus onset. We then define a baseline by the first 7 time bins (about 14 ms) and subtracted it from the RMS curves.

Because the electrophysiological dataset was mostly collected from the left hemisphere of the mouse brain, only the data of right-stimulus trials captures the early sensory response (within the first 100ms after stimulus onset, see **Fig.** S11), and we expect to get similar responses for the left-stimulus trials if we had sufficient recordings from the right hemisphere. Therefore, we take the average stimulus response to right-stimulus trials as the data curves in **Fig.** 6b, assuming the left-stimulus trials to have the same responses. In the model, the stimulus response curves were based on the difference between right and left stimulus response units (*x*_*S, model*_(*t*; *c*) = ⟨*s*_*R*_(*t*) − *s*_*L*_(*t*) ⟩), averaged for right-stimulus trials of each contrast within the first 100 ms following stimulus onset. The total loss used for optimization combines a sum of squared errors (SSE) term and a squared deviation term between the empirical and model-predicted SNR values across all contrasts:

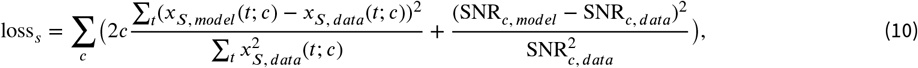

where the SSE is normalized by the total power in the data and scaled by the contrast value to weigh high contrast responses more, and the SNR term is normalized by the squared value of data SNR. At this fitting stage the prior modulation of gains and biases (*g*_*i*_, *δ*_*i*_) are set to zero because the curves being fit sum over both block sides, thus prior modulation effects should largely cancel. This simple model generates reasonable contrast-dependent stimulus responses, **Fig.** 6b (*R*^2^ = 0.872), with the parameter values of *w*_*ss*_ = 0, *w*_*as*_ = 28, *τ*_*a*_ = 223, *α* = 1.6, *β* = 0.16, *σ* = 35, *λ* = 2.

The second stage of fitting process targets all the remaining free parameters for integrator/choice *C*, movement *M*, and prior *P* dynamics: weights *w*_*cc*_, *w*_*mm*_, *w*_*pp*_, *w*_*cs*_, *w*_*mc*_, *w*_*pc*_, prior gain and bias modulations *g*_*i*_, *g*_*m*_, *δ*_*i*_, *δ*_*m*_, and action thresholds *θ*. We restrict *w*_*pp*_ in a tight range [0.495, 0.4999] so that **p** is well-tuned based on the observed long time-constant of prior integration spanning many trials, and attempt to match the model’s responses and the prior distance curves 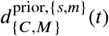 to the neural responses and prior distances. We calculate the RMS activity from integration/choice regions and movement regions (**Fig.** 2), 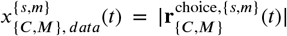 for choice left and right trials aligned to stimulus and movement onset, baseline-corrected by shifting the curves to start at 0 at stimulus onset, and truncated to the 150 ms after stimulus onset or before movement onset. We exclude the initial 15 time points (about 30 ms) after stimulus onset from the loss function to remove contribution from the transient onset response observed in data. The model’s C and M responses for right and left choice trials (*x*_{*C,M*}, *model*_(*t*) = ⟨{*c, m*}_*R*_(*t*) − {*c, m*}_*L*_(*t*) ⟩ for right choice, ⟨{*c, m*}_*L*_(*t*) − {*c, m*}_*R*_(*t*) ⟩ for left choice) are compared to these data curves with the same alignment, and we calculate a normalized SSE loss between them. To ensure the model’s activities can return to the 0 baseline before the start of each trial, we penalize the choice/integrator and movement neurons’ nonzero activities during [-400, -100] ms before stimulus onset (inter-trial window, but before prior biases are applied).

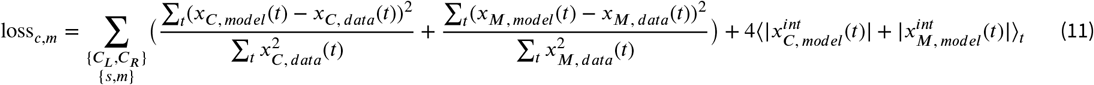

Additional regularization terms enforced that the mean prior representation in the model tracked the block side correctly, namely that 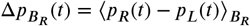 for right blocks and 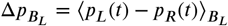 for left blocks should both have positive mean values over time (we selected the 150 time steps (about 300 ms) before the stimulus onset for this calculation). We also penalized the difference between the 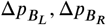 mean values, normalized by the sum of their squared means.

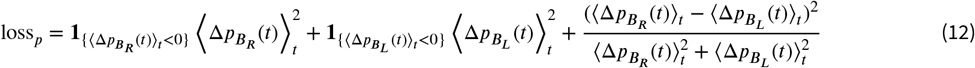

The prior distance data curves for integration/choice 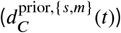 and movement initiation areas 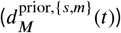 are shifted by subtracting the mean of shuffled control distance curves at each time bin. We calculated the model’s prior distance curves in the same way as in data (restricted to the same choice and stimulus sides and sum over them; with the same stimulus and movement onset alignment), and calculated the SSE normalized by total power in the data curves in the loss term.

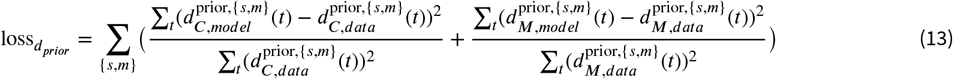

Overall, the second stage of fitting combined the loss terms above, 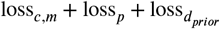 for optimization. We obtain the following parameter values: *w*_*cc*_ = 0.43, *w*_*mm*_ = 0.27, *w*_*pp*_ = 0.496, *w*_*cs*_ = 0.17, *w*_*mc*_ = 0. 50, *w*_*pc*_ = 1.6 × 10^−5^, *g*_*c*_ = 987, *g*_*m*_ = 6, *δ*_*i*_ = 21, *δ*_*m*_ = 10^−3^, and *θ*_*d*_ = [0.76, 0.40] for concordant/discordant trials. For the alternative model with direct bias (Eq.9), the fitting process produced the following parameter values: *w*_*cc*_ = 0.43, *w*_*mm*_ = 0.27, *w*_*pp*_ = 0.496, *w*_*cs*_ = 0.17, *w*_*mc*_ = 0.50, *w*_*pc*_ = 1.6 × 10^−5^, *g*_*c*_ = 932, *g*_*m*_ = 6, *δ*_*c*_ = 80, *δ*_*m*_ = 10^−3^, and *θ*_*d*_ = [0.76, 0.41] for concordant/discordant trials.

## Supplementary Material

**Figure S1.**
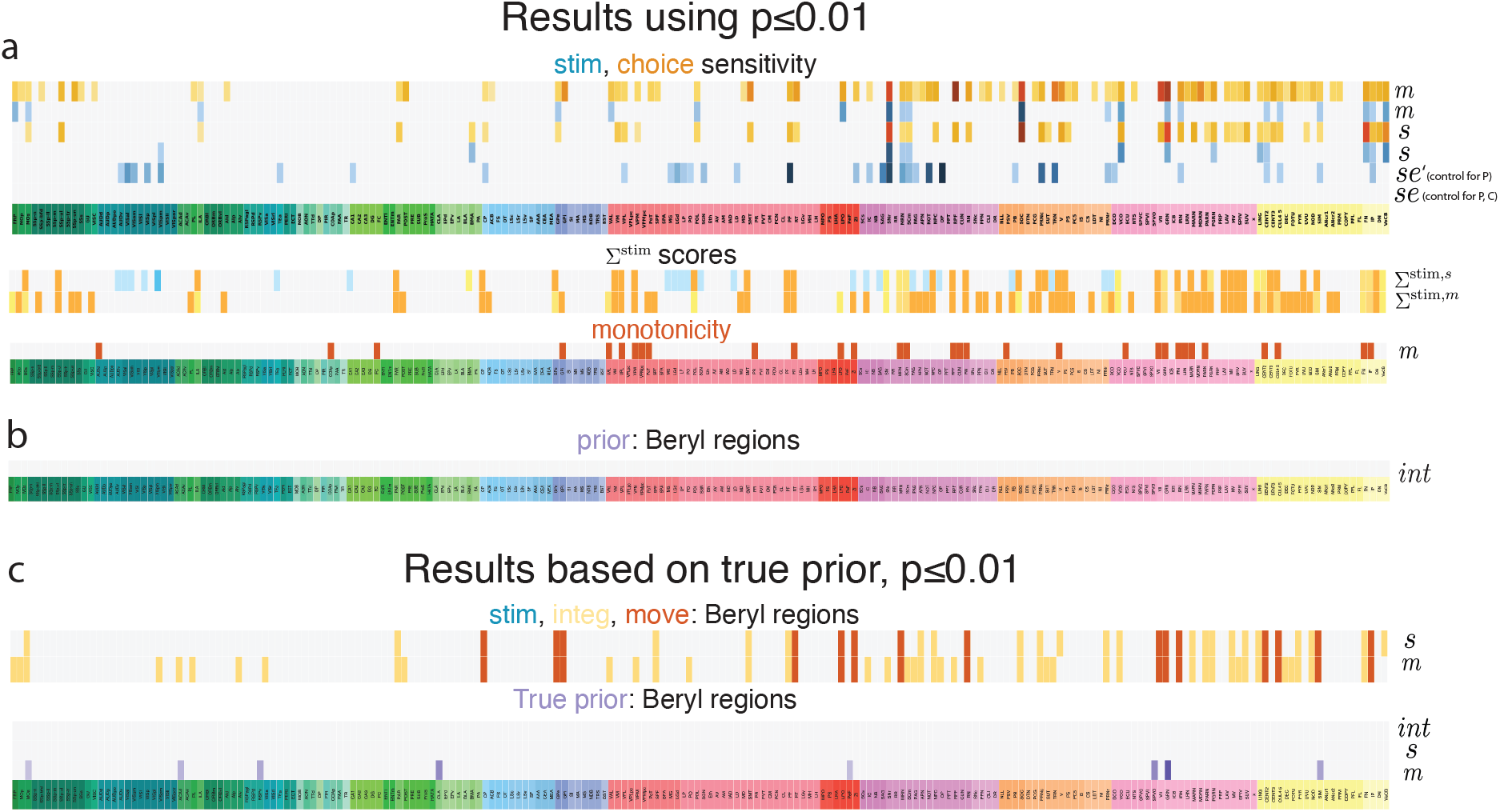
Identification of sensory response, integration, movement initiation, and prior regions, using *p* ≤ 0.01. **a** Top table: stimulus (rows with blue cells) and choice sensitivity (rows with orange cells). Darker colors indicate larger effect size 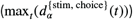: stimulus-onset aligned, 0-150 ms; *se*: stimulus-onset aligned early stimulus period, 0-80 ms, controlling for correlations with prior and choice sides; *se*^′^: stimulus-onset aligned early stimulus period, 0-80 ms, only controlling for correlations with prior side; *m*: movement-onset aligned, -150-0 ms. Middle and bottom tables: 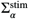 metric (darker blue: 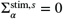 using 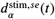; lighter blue: 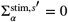 using 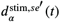; darker orange: 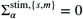; other colors represent values in between), and monotonicity metric. Both metrics are defined in Results. Light gray cells: not significant (top table) or undefined (middle and bottom tables) due to lack of significant stimulus or choice sensitivity. **b** Summary table of prior sensitivity during the inter-trial (*int*) window. **c** Results using the true block prior (in contrast to the subjective prior presented in the main results and in a-b). The first table shows the sensory response, integrator, and movement initiation regions using the true block prior when controlling for the three-way dichotomies of stimulus, choice, and prior (details in Methods), and the second table summarizes prior encoding regions during inter-trial, stimulus, and movement periods.

### Prior in the inter-trial window

We briefly examine – for continuity with recent work – the encoding of the prior in the inter-trial period (here referring to the [−400, −100] ms interval aligned to stimulus onset and designated *int*) ^2^, using the subjective action kernel prior. No stimulus was presented or choice made in this interval, therefore we do not condition on stimulus or choice side:

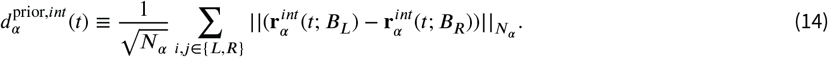

We find very sparse neural coding of inter-trial prior (**Fig.** S1b,S2b), with 5/208 significant prior encoding regions using *p* ≤ 0.05 and 0 significant prior regions using *p* ≤ 0.01. Correlating neural responses with true block prior reveals no significant inter-trial prior-coding regions (**Fig.** S1c, S1e) because the animal is not privy to the true block side, consistent with past findings ^1,2^. We consistently find sparser coding of prior during trials using the true block than 3hen using subjective estimates (**Fig.** S1c, S1e, S3).

### Prior effects on sensory areas

We examine the sensory regions under less stringent inclusion criteria for stimulus side (only controlling for stimulus-prior correlation, Eq.3) and find three of them that show significant prior side sensitivity, *p* ≤ 0.01 (intersection of the 1st row light blue regions and 2nd row purple regions in the table in **Fig.** 4a). Among these three regions, SAG has a reasonable stimulus sensitivity curve that shows significant response within the first 40 to 80 ms after stimulus onset (**Fig.** S4a top left), but its prior sensitivity is not significant during the same time window, indicating that prior is not modulating the sensory response in SAG. The other two regions, PAG and DCO, do not show a typical stimulus response trace that starts insignificant at stimulus onset and ramps up around 40 ms after stimulus onset. Instead, they show a significant initial offset in stimulus sensitivity similar to prior sensitivity, and without a significant gain response later on (gain *p* > 0.01 for both regions), suggesting their significant stimulus sensitivity is a result of biases from previous trials. We also examined the prior sensitivity in other light-blue stimulus response regions that are known to be relevant to stimulus processing from previous work ^1^, such as LGd, VISp, and VISpm (**Fig.** S4b), and they clearly do not show significant prior coding. Therefore, we conclude that even with the expanded set of stimulus processing regions using less stringent controls, there is still no significant prior modulation on sensory processing in the current task.

**Figure S2.**
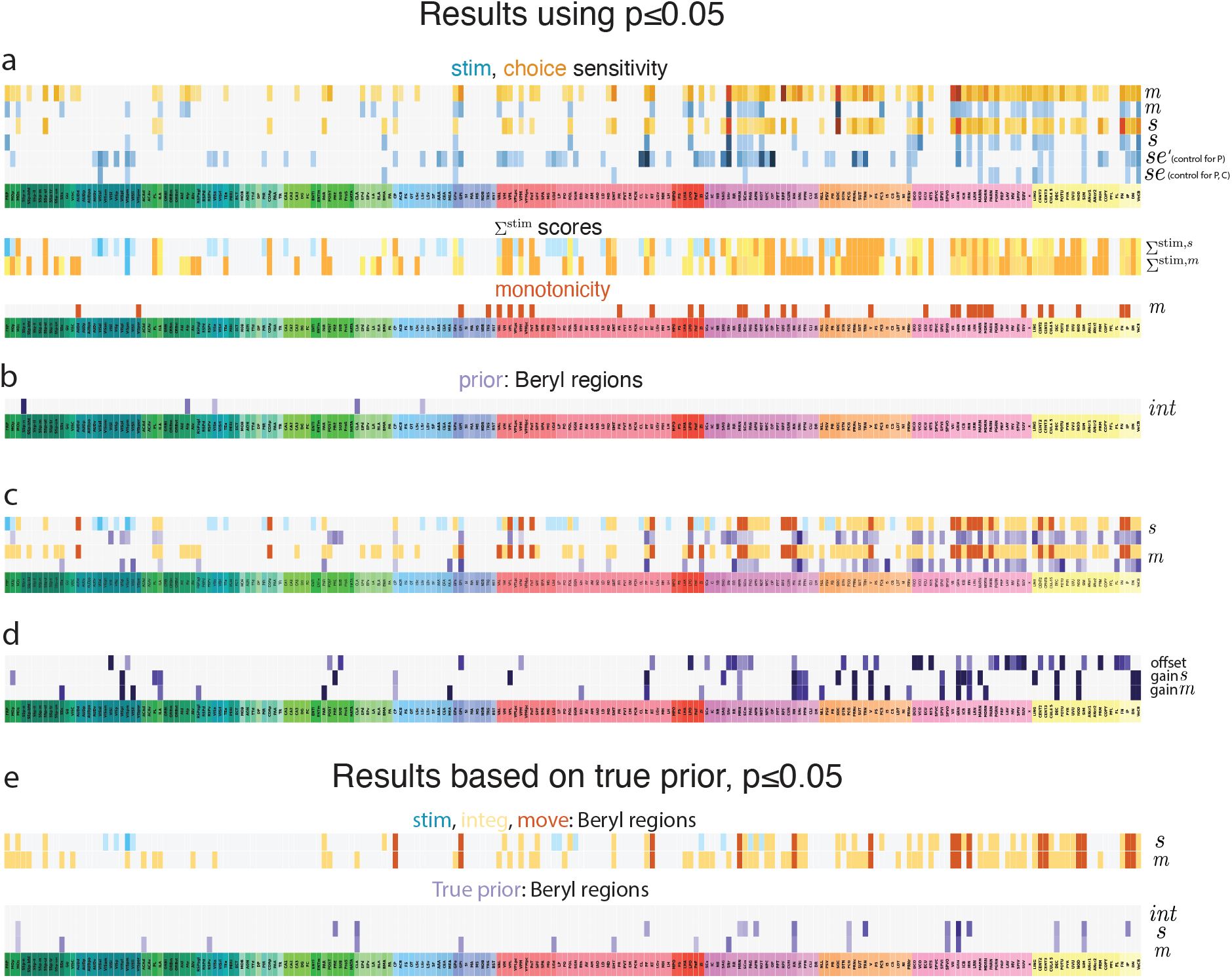
Identification of sensory response, integration, movement initiation, and prior regions, using *p* ≤ 0.05. **a** Top table: stimulus and choice sensitivity. Middle and bottom tables: Σ^stim^ metric, and monotonicity metric. **b** Summary table of prior sensitivity during the inter-trial window. **c** Summary table of region functionality and prior sensitivity when aligned to stimulus and movement onset, similar to **Fig.** 4a. **d** Summary table of prior offset effect, gain effect aligned to stimulus onset, and gain effect aligned to movement onset, similar to **Fig.** 4h. **e** Results using the true block prior (in contrast to the subjective prior presented in the main results and in a-d). The first table shows the sensory response, integrator, and movement initiation regions, and the second table summarizes prior encoding regions during inter-trial, stimulus, and movement periods.

**Figure S3.**
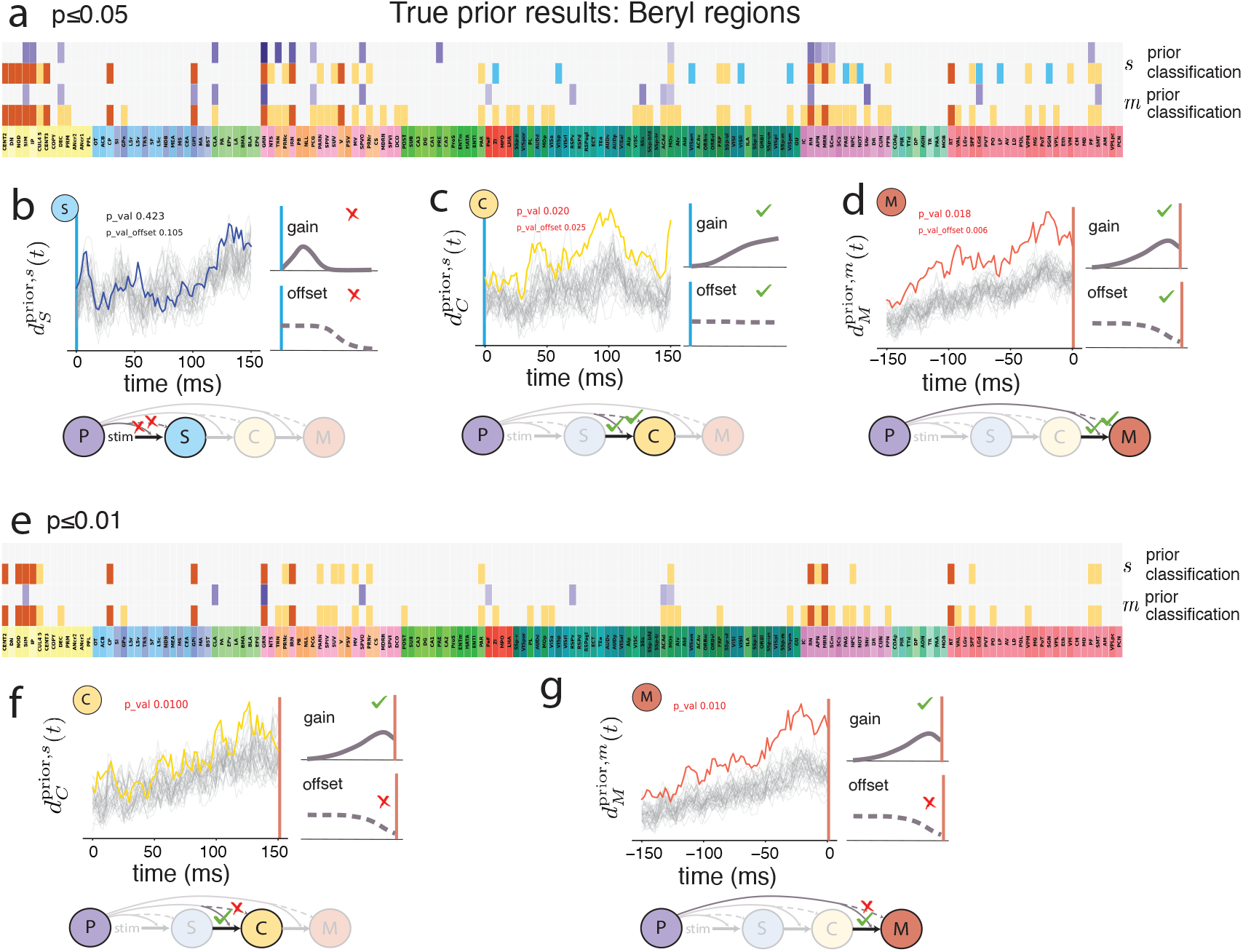
Prior effects on Beryl regions using the true block prior (two different significance thresholds). We present results for the same analyses as in Fig.4 but using the true prior instead of subjective prior. **a-d** are results using *p* ≤ 0.05 for significance tests, and **e-g** are using *p* ≤ 0.01. We reproduce the main results of prior modulation on integrator and movement initiation regions using the true prior with *p* ≤ 0.05, but the direct offset modulations are weaker than gain modulations and are not significant using *p* ≤ 0.01. Also, purely sensory response regions have relatively weaker signals and are only identified using *p* ≤ 0.05.

**Figure S4.**
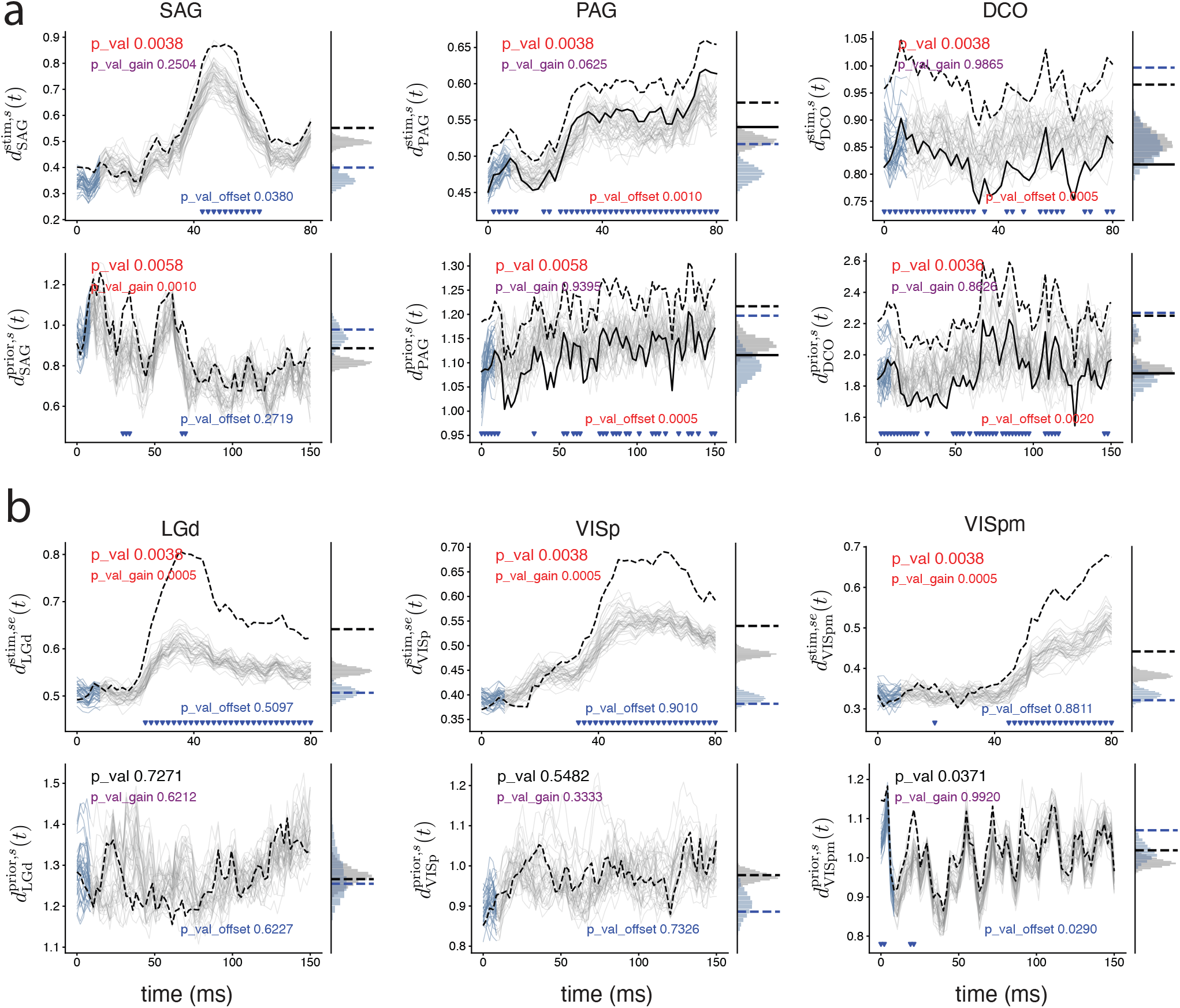
Prior effects on sensory areas. **a** Stimulus (top row) and prior (bottom row) sensitivity (euclidean distance as defined in Eq.1,6) in SAG, PAG, and DCO, where both stimulus and prior coding are significant. Dashed black curves: raw euclidean distance over time (main panel) and its time-average (right panel); blue dashed curve in the right panel: time-average of the raw euclidean distance over the first five time bins, where we characterize the effect and significance of initial offsets; solid black curve: euclidean distance over time subtracting the effective offset, if offset is significant (main panel), and its time-average (right panel). Gray curves in main panel: shuffle controls, with their first five time bins colored in blue; gray histogram in the right panel: distribution of the gray shuffle curves’ time-average; blue histogram: distribution of the shuffles’ time-average over the first five time bins. The triangles at the bottom indicate significance in the time bins. **b** Stimulus (top row) and prior (bottom row) sensitivity in example regions where only stimulus coding is significant. **c**Prior effects on Beryl regions using the subjective action kernel prior, *p* ≤ 0.05.

**Figure S5.**
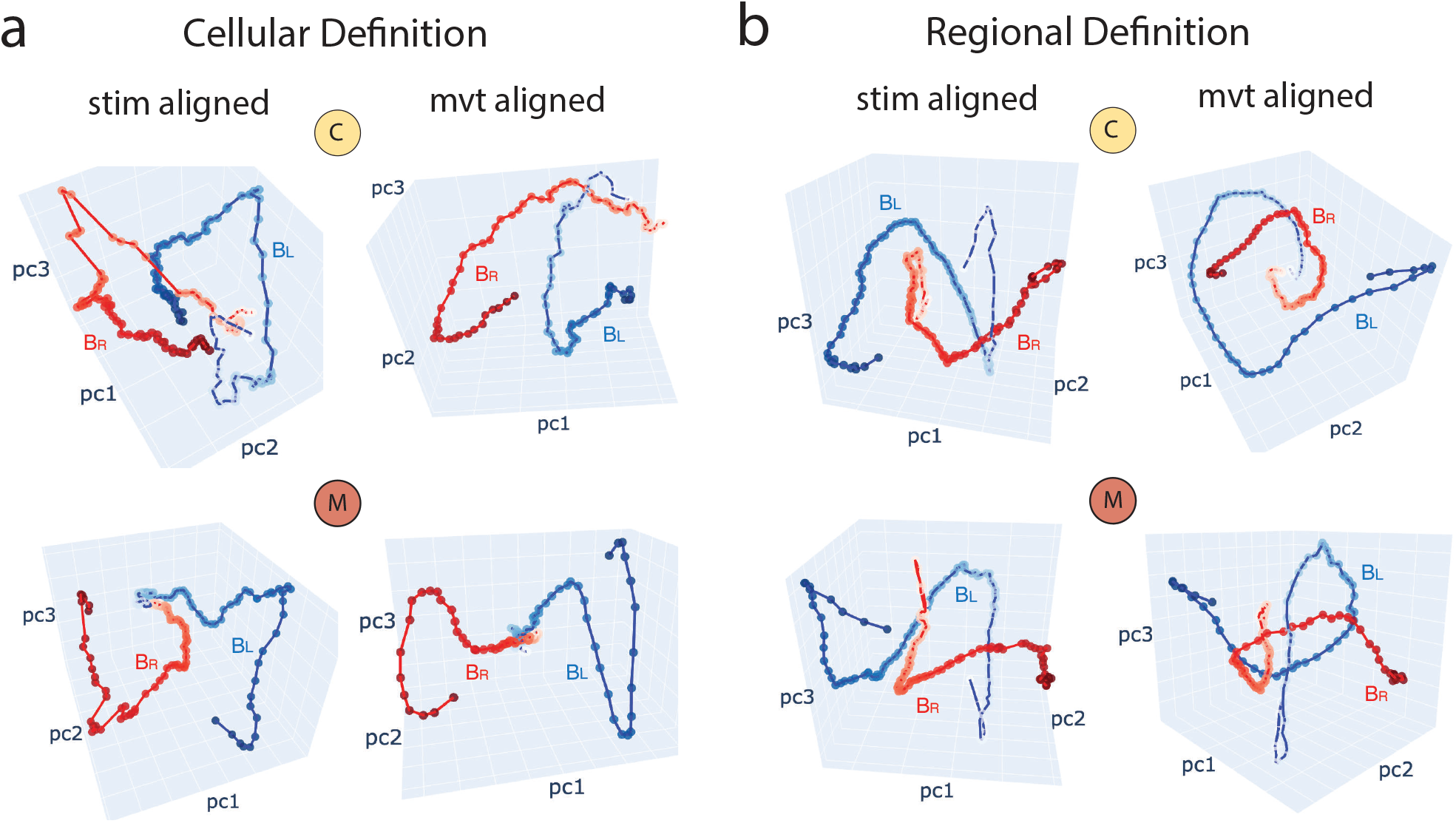
Trajectories by prior side projected onto dimensionally reduced state space. We applied Principle Component Analysis (PCA) to the original high-dimensional prior L, R trajectories and plot them in the 3-D space defined by top three principle components. We show the right stimulus, right choice trajectories 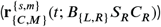 as examples, and other trajectories are quantitatively similar. **a** Results on cellular-level definition of integrator and movement initiation, aligned to stimulus onset (0 − 150ms after) and movement onset (0 − 150ms before). Darker color indicates later times in trial. **b** Results on regional-level definition, combining all regions identified as integrator/ movement regions that show significant prior modulation during trial.

**Figure S6.**
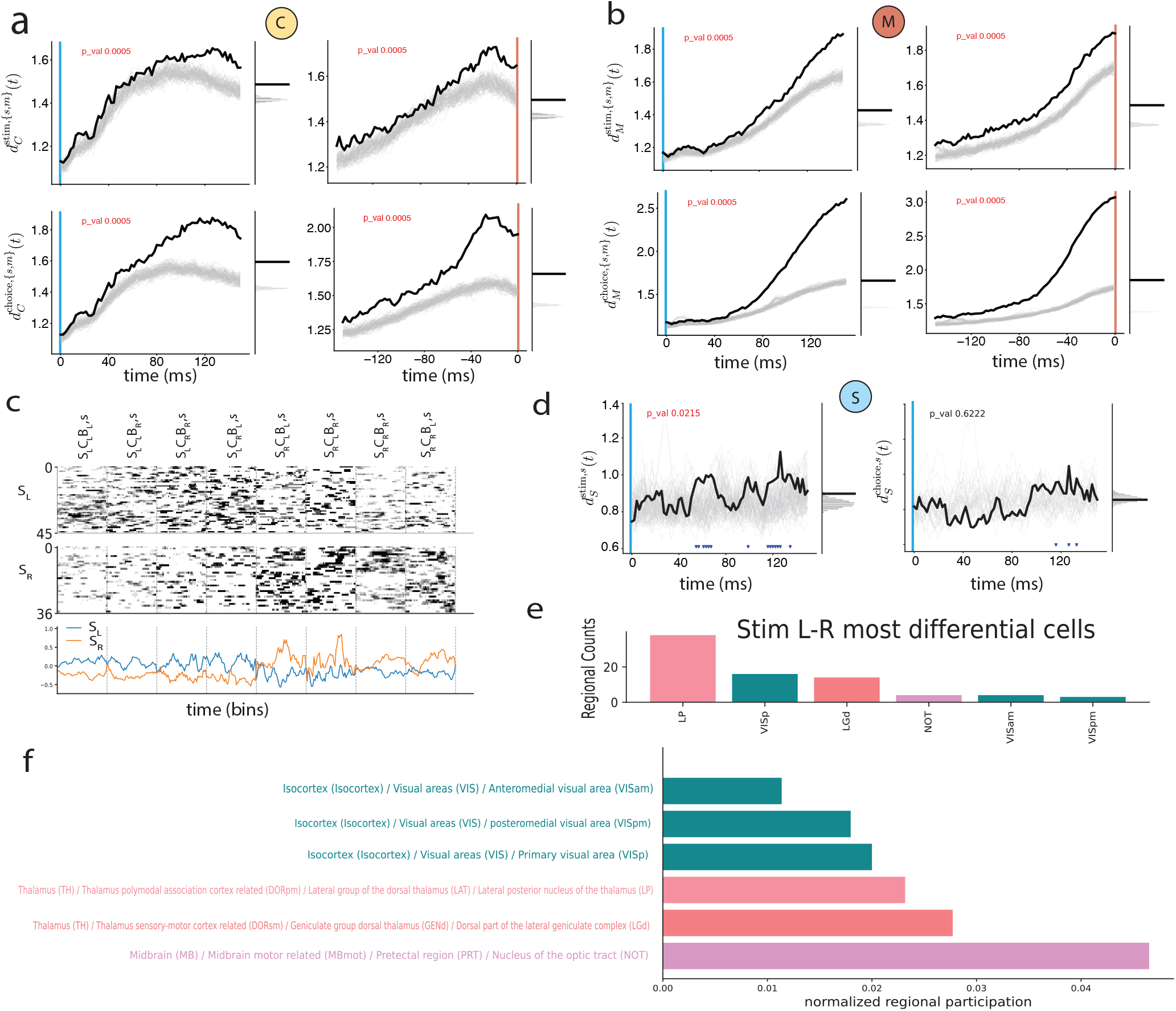
Stimulus and choice sensitivity of region-independent cellular response groups. **a** The integrator cells defined by Rastermap algorithm show significant stimulus (top row) and choice (second row) sensitivity for both stimulus aligned (left column) and movement aligned (right column) time windows. **b** Similarly, the movement initiation cells show significant stimulus and choice sensitivity when aligned to stimulus onset and movement onset. **c** The stimulus response cells are defined from sorting cells with the most early stimulus response and the least choice response in relevant Beryl regions. Top two panels: raster plots of top stimulus L and R responding cells, sorted by the Rastermap algorithm. Bottom panel: average PETH of these top stimulus L and R responding cells. **d** The stimulus response cells show a significant response stimulus difference (left) but not choice difference (right). **e-f** Regional distribution of cells in the stimulus response group. **e** shows a histogram of cell counts, **f** shows the same data normalized by total number of cells in the regions, as well as the full names and anatomical hierarchy of these regions.

**Figure S7.**
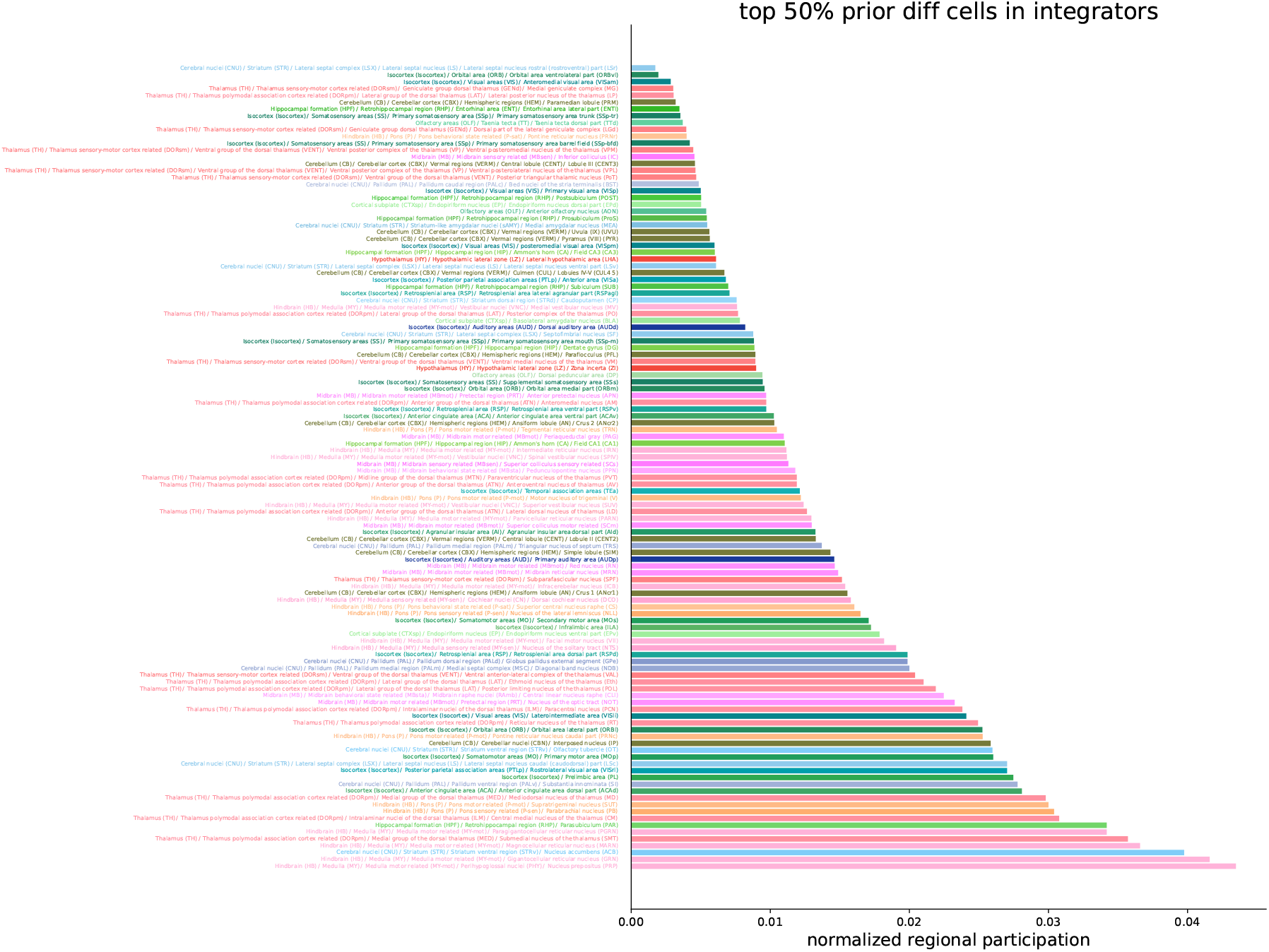
Top prior differentiating cells: integrator. We sort the integrator cells by their average block L-R response difference, and show the regional distribution of the top 50% of these cells. For each region we list their full names and anatomical hierarchy.

**Figure S8.**
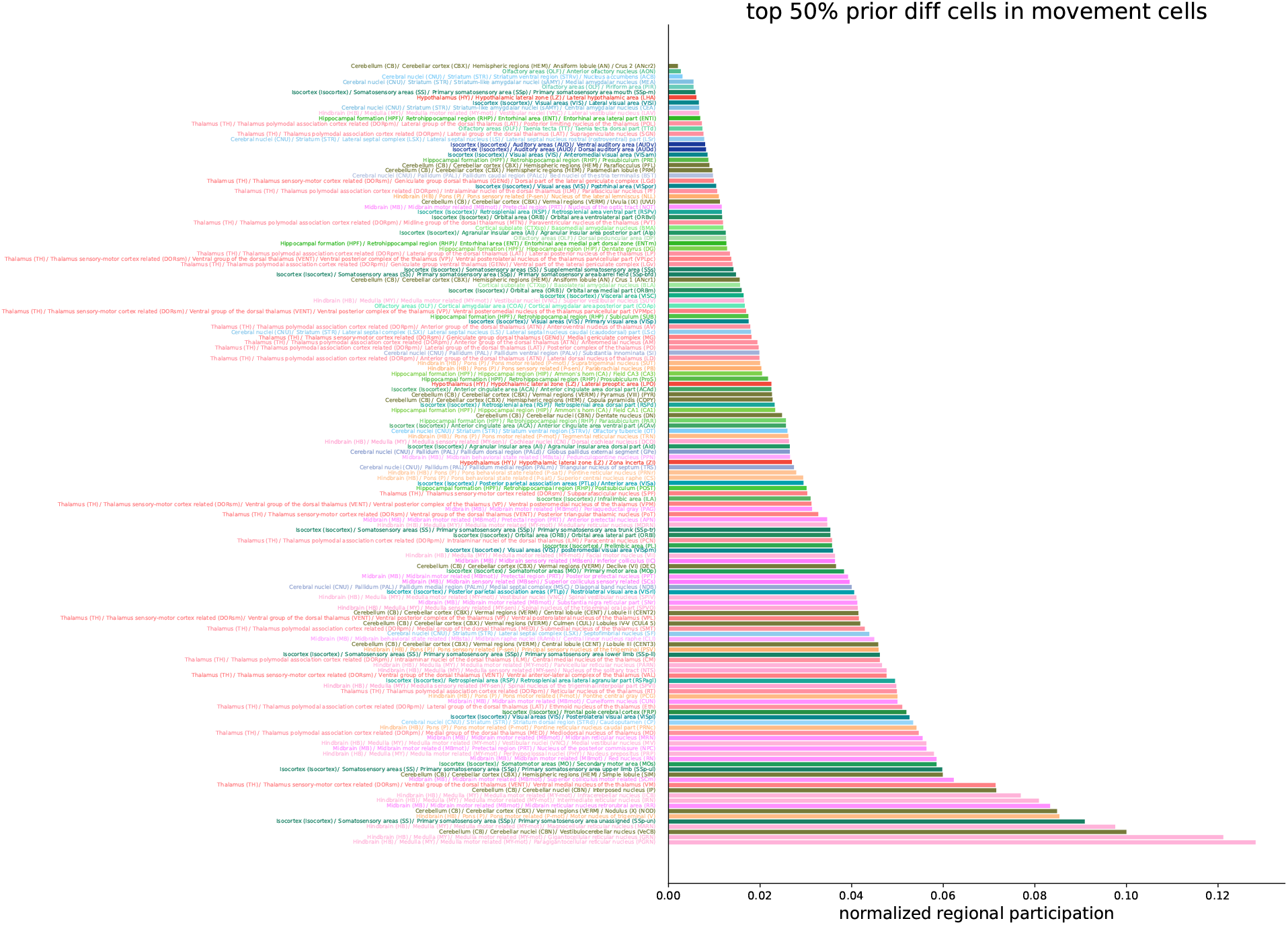
Top prior differentiating cells: movement. We sort the movement cells by their average block L-R response difference, and show the regional distribution of the top 50% of these cells. For each region we list their full names and anatomical hierarchy.

**Figure S9.**
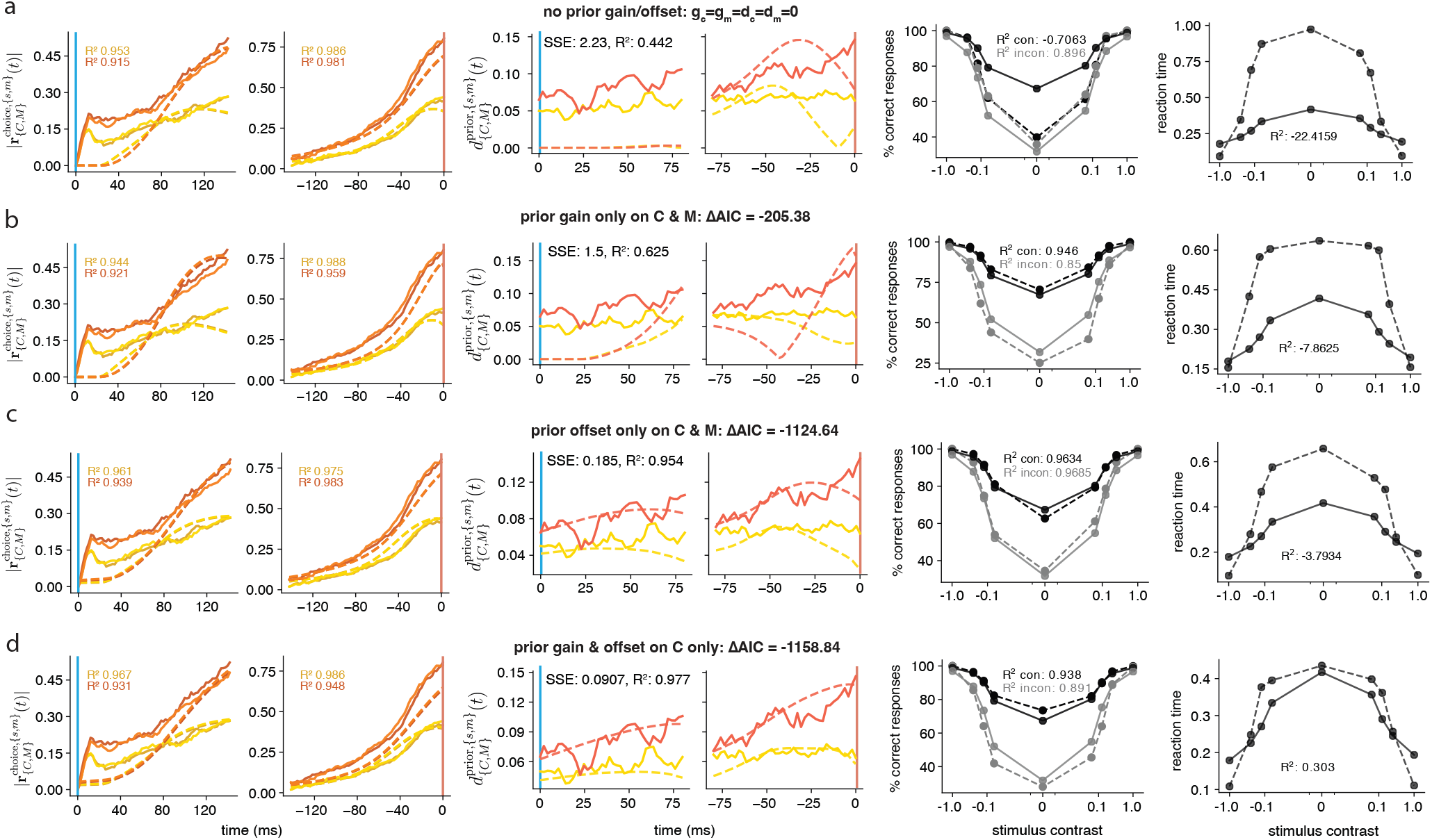
Different models of prior modulation, using offset as an integrated input. We fit to the data four alternative models using Eq.7: **a** with no prior modulation, **b** only gain modulation on the choice/integration and movement nodes, **c** only offset modulation on the choice/integration and movement nodes, and **d** gain and offset modulations on only the choice/integration nodes. The fitting process to choice and movement response curves and prior distance curves are shown in the left and middle columns, and the resulting model-predicted behaviors are shown on the right. Dashed: model; solid: data.

**Figure S10.**
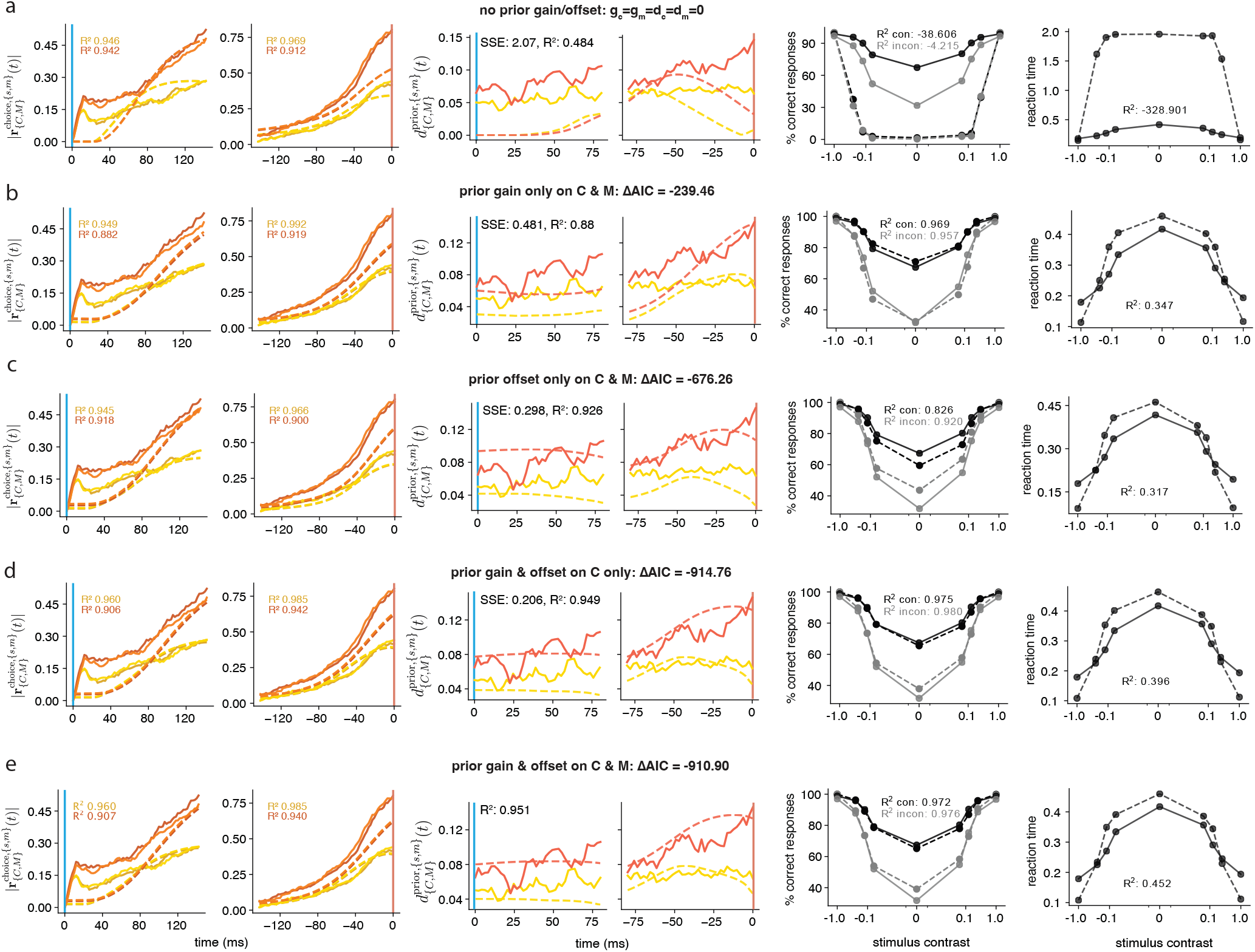
Different models of prior modulation, using direct offset bias. We fit five models to the data using Eq.9: **a** with no prior modulation, **b** only gain modulation on the choice/integration and movement nodes, **c** only offset modulation on the choice/integration and movement nodes, **d** gain and offset modulations on only the choice/integration nodes, and **e** with gain and offset modulations on both choice/integration and movement nodes. The fitting process to choice and movement response curves and prior distance curves are shown in the left and middle columns, and the resulting model-predicted behaviors are shown on the right. Dashed: model; solid: data.

**Figure S11.**
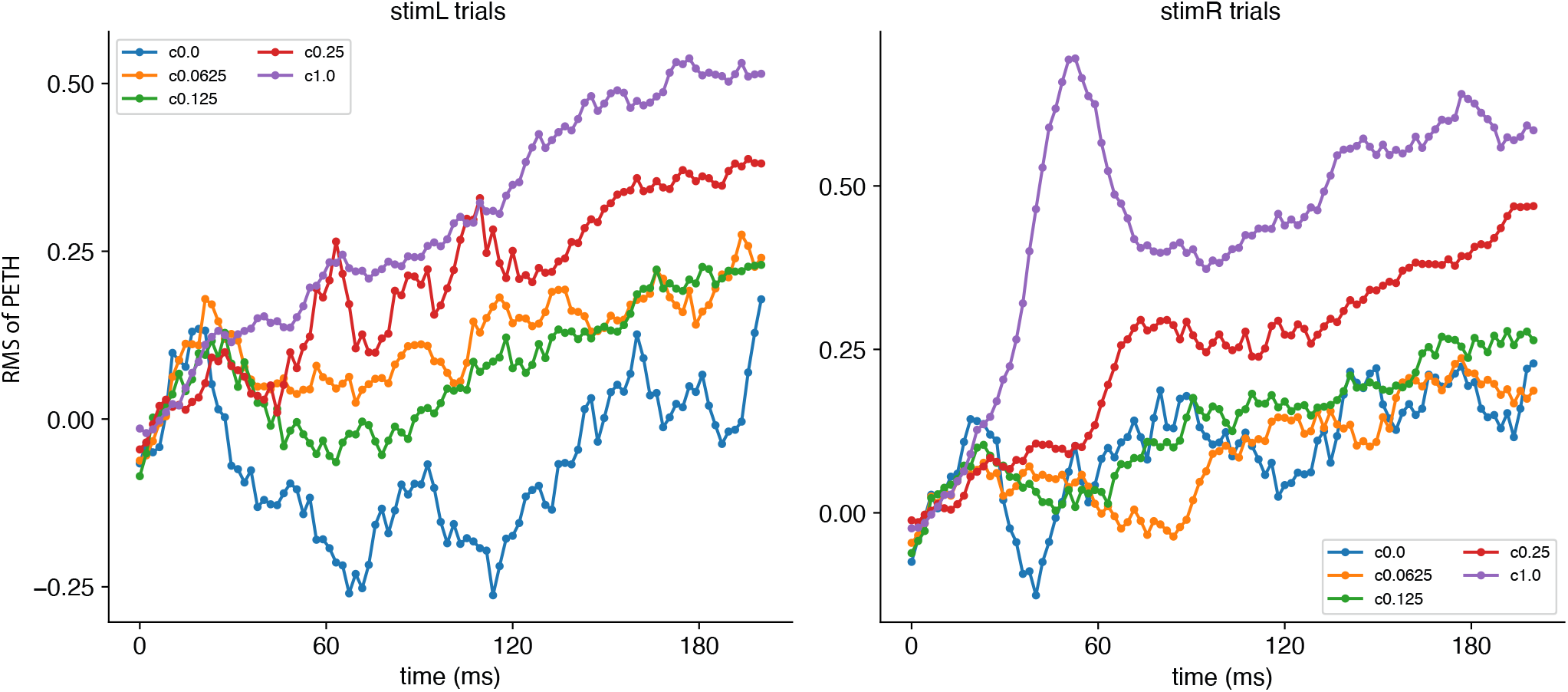
Mean sensory response to left and right stimulus. We plot the root-mean-square response of neurons in sensory areas (VISpm, VISam, FRP, VISp, VISli, LGd, LP, NOT) to left and right stimulus trials by contrast. Time=0 corresponds to stimulus onset time.

